# *Delta-Marches:* Generative AI based image synthesis to decode disease-driving morphologic transformations

**DOI:** 10.1101/2025.03.18.643999

**Authors:** Thuong Nguyen, Vandana Panwar, Vipul Jarmale, Averi Perny, Cecilia Dusek, Qi Cai, Payal Kapur, Gaudenz Danuser, Satwik Rajaram

**Affiliations:** Lyda Hill Department of Bioinformatics, University of Texas Southwestern Medical Center, Dallas, TX, USA; Department of Pathology, University of Texas Southwestern Medical Center, Dallas, TX, USA; Kidney Cancer Program, Simmons Comprehensive Cancer Center, University of Texas Southwestern Medical Center, Dallas, TX, USA; Department of Urology, University of Texas Southwestern Medical Center at Dallas, Dallas, TX, USA; Institute of Human Biology, Roche Pharma Research & Early Development of F. Hoffmann-La Roche Ltd, Basel, Switzerland

## Abstract

Deep learning reveals that tissue morphology contains rich pathophysiological information beyond human understanding. However, approaches to convert these spatially distributed signals into subcellular insights informing disease mechanisms are lacking. We introduce Delta-Marches, an interpretability-first approach that nominates distinguishing morphological features rather than explaining existing models’ decisions. Delta-Marches simulates idealized morphological changes between classes by coupling latent-space traversals with generative AI. Comparing each image to its class-shifted counterpart allows feature extractors to infer the most affected aspects, reducing sample-to-sample variability and resolving transformations at subcellular resolution. Prototyped in renal carcinoma grading, Delta-Marches generates realistic grade transitions and pinpoints tumor-cell nuclear phenotypes as key determinants. It also reveals reduced vasculature with increasing grade, a known pattern absent from standard rubrics. Applied to dysplastic progression in colorectal tissue, it recovers glandular remodeling and goblet-cell loss, generalizing across tissue types and scales. These results show Delta-Marches parses complex image phenotypes and catalyzes hypothesis generation.

## INTRODUCTION

The ultimate opportunity of applying artificial intelligence (AI) to biomedicine (*1, 2*) is to exploit subtle and complex patterns in data with a sensitivity and precision that surpass human intuition—and to transform these patterns into meaningful scientific and clinical insights. In genomics, for example, models (*3–5*) have predicted regulatory motifs and revealed syntax rules, such as the spacing and orientation of transcription factors binding sites, that govern enhancer activity, yielding insights that extend well beyond traditional motif discovery.

In contrast, tissue microscopy, while central to clinical workflows, remains underutilized for AI-driven discovery, despite widespread recognition that these images contain rich, untapped insight into disease mechanisms (*6, 7*). This gap is evident in tumor histopathology. Historically, the rules of tumor assessment have been developed under criteria of ensuring consistency across observers rather than capturing the full range of morphologically predictive information. Many disease-driving features involve subtle molecular or spatial patterns that interact in complex, non-linear ways, making them difficult to isolate by visual inspection. Classical machine learning approaches (Fig. 1A top), which rely on manually engineered features, were initially expected to offer more comprehensive diagnostics with a high-level of interpretability. In practice, however, even generating relevant biological features, such as identifying cell types, can be a substantial undertaking (*8–19*). Moreover, sample-to-sample variation often masks class-defining morphological differences, and without counterfactual examples—samples that differ only in the property of interest—the true discriminative features remain difficult to pinpoint. Modern deep learning has demonstrated remarkable performance across a range of pathology tasks, achieving expert-level tumor grading and subtyping and even surpassing human intuition in tasks such as predicting mutation status directly from morphology (*20–41*). Despite this success, these models have rarely translated predictive power into biological insight.

**Fig. 1.**
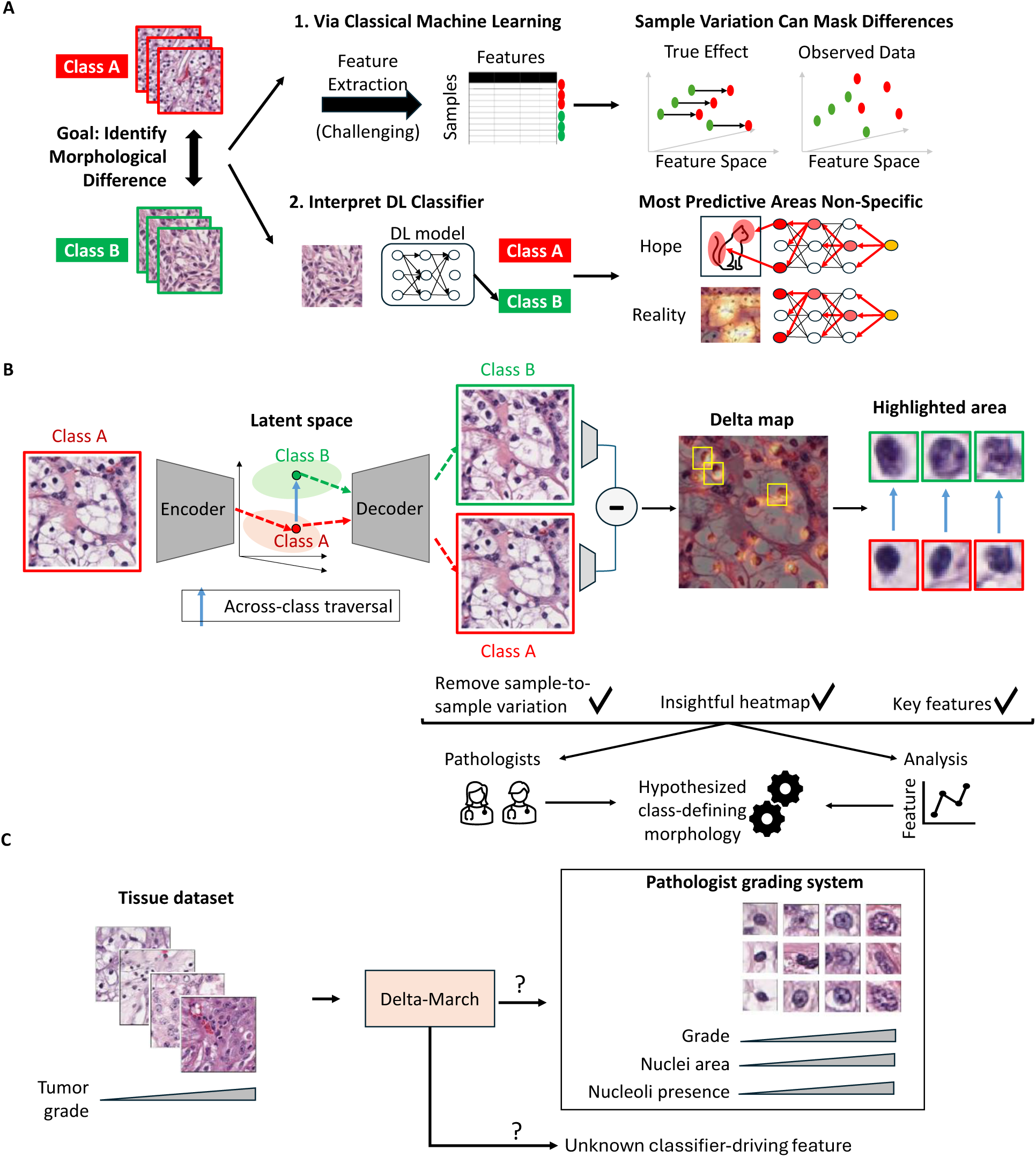
Overview of the study. **(A)** Understanding of the association between tissue morphology and its biology has evolved through centuries of observation and empirical study. In recent years, rule-building by human inspection has been complemented by classical machine learning (ML) techniques that can analyze features across biological categories by rigorous statistics. However, tissue complexity and within- and across-tumor variability hinders the engineering and interpretation of meaningful features. Deep learning (DL) excels in the classification of disease states by tapping into complex image features that are inaccessible to humans. However, DL models are seen as “black boxes” with limited transparency on which image information drives the classification. Efforts in developing interpretable DL (IDL) models have attempted to overcome this deficit by revealing the links between input data and predictions. However, traditional IDL methods tend to rely on saliency maps that mark classifier-informing regions in broad strokes but fail to extract from histopathological data the local and often spatially dispersed image properties that are critical for the classifier performance. **(B)** The framework proposed in this work implements an autoencoder model and latent space traversal (blue arrow) to generate synthetic images of the same tissue with different classes. These synthetic images are utilized to create a Delta-Map, which underscores substantial across-class transformations and provides insights into the mechanisms driving morphological changes associated with biological progression. **(C)** Primary benchmark for the present study: Evaluate the ability of the proposed model to autonomously learn clinically-established tumor grading rules and in potentially creating new rules that may inform an enhanced tumor grading procedure.

While there is much work in interpretable deep learning (IDL) and Explainable AI (xAI), much of it focuses on learning how a classifier makes its decisions (*42–52*) rather than revealing biological insights. As standard models are typically optimized for classification prowess rather than post-hoc deconstruction, this creates a gap between these objectives. This limitation is exacerbated in tissue microscopy. Unlike natural images where a class is often localized to a single object, pathological signals are spatially scattered and intermixed (Fig. 1A bottom). Because discriminative convolutional neural networks (CNNs) aggregate these diffuse signals into a single class probability, methods that retroactively explain this process (*53–57*) inevitably smear fine-grained details into broad, nonspecific heatmaps. This tradeoff persists with more recent architectures such as Vision Transformers (ViTs), albeit with a distinct failure mode. Driven by the objective of finding the most efficient path to a classification decision, ViTs often spotlight a minimal set of highly discriminative tokens (*58, 59*), such as a single tumor nucleus (Fig. S1). For a human observer, this sparsity is confusing: it leaves biologically identical tumor nuclei unhighlighted, appearing as false negatives undermining the corresponding biological hypothesis. Thus, while traditional xAI is important to establish confidence in existing models, it remains ill-suited for generating novel biological hypotheses in microscopy.

We previously developed a conceptually distinct IDL approach leveraging generative AI —not to explain a DL prediction in the traditional sense, but to explore the morphological basis of an image-based classifier of metastatic propensity of single melanoma cells in culture (*60*). Specifically, we trained a latent representation capable of generating images of cells with different levels of metastatic spreading. By examining the warping of images as we traversed the latent space between areas of high and low metastatic propensity, we identified extension of pseudopods as the defining feature of high metastatic potential. Both the classifier and the IDL-predicted morphological behavior of aggressively metastasizing melanoma cells were validated in follow-up studies using human xenografts in mice (*60, 61*). Of note, it would not only be very difficult to human-engineer a detector of pseudopodal extension in these images, but the cell-to-cell heterogeneity completely obfuscates the differential in pseudopodia extension as a defining characteristic of low vs. highly metastatic cells. Rather, this discovery was made possible by the power of generative AI to gradually parse the complex mix of phenotypes across cells with differing metastatic potential even beyond the range of available experimental data. While several studies (*62–64*) have since applied generative AI models to histopathology for expanding datasets and visualizing class transformations (*65–73*), these approaches primarily function as image synthesis tools rather than analytical engines. By producing synthetic transformations with either coarse or implicit characterizations, current methods effectively shift the interpretability burden back to the human observer, who must subjectively ‘spot the difference’ to resolve subtle, subcellular features amidst complex, heterogeneous tissue architectures. Consequently, the potential of generative AI to elucidate the relationship between tissue structure and disease state has remained largely untapped.

Here, we design a hypothesis generation framework that extends the concept of interpretability via latent space traversal from single cells to patient tissue samples through two key innovations (Fig. 1B). First, we adopt diffusion models (DMs) (*74–79*) to learn a meaningful high level semantic latent space that is agnostic to interchangeable low level image features, which are nonetheless crucial for the production of realistic images (Fig. 1B). Second, we introduce the concept of the *Delta-March*, a system to analyze changes between counterfactual images, images of the same tissue that differ in class, produced by the latent space traversal to identify the disease-associated features. By generating estimates of counterfactual differences with sub-cellular resolution, our method enables deep quantitative analysis of morphological transformations, robustly capturing subtle and spatially dispersed patterns and providing subcellular-resolution interpretation even in complex tissue architectures. As a test case, we demonstrate that the proposed model faithfully recovers the clinically applied grading rules for clear cell renal cell carcinoma (ccRCC). Delta-Marches also highlight an additional feature that, while accepted by pathologists as plausible, has yet to be entered into the official grading scheme. Beyond the primary ccRCC grading task, we evaluated the framework’s ability to generate trajectories consistent with the evolutionary relationships among a patient’s tumor regions, using a multi-regional sequencing dataset. Finally, to test its ability to scale across systems and physical length scales, we applied it to the normal-to-dysplastic transformation in colorectal tissue. This provides a glimpse of the potential of this AI framework to complement existing diagnostic approaches with novel biomarkers and to assist clinicians in the generation of hypotheses on cancer biology that has so far escaped the current body of medical knowledge.

## RESULTS

### Tumor grading rules as a benchmark for hypothesis generating IDL

Pathologists classify ccRCC into four grades of tumor aggressiveness based on hematoxylin and eosin (H&E) stained histopathology slides. The scoring systems have been updated over time but are largely focused on nuclear size and nucleolar prominence, both of which increase with higher-grade aggressive tumors (*80–82*). Specifically, distinctions of grades 1, 2, and 3 are purely based on these features, while grade 4 also considers non-nuclear properties of tumor cells such as sarcomatous or rhabdoid differentiation, which are associated with poor prognosis and more aggressive tumor behavior (*80–82*). Additionally, while not part of the official grading rules, the extent of vasculature is known to decrease with increasing grade (*83–85*). We note that ccRCC tumors exhibit a high degree of intra-tumor heterogeneity (*20*), and grade is traditionally assigned at a patient level based on the most aggressive area within a tumor. Consequently, conventional datasets, and models derived from them, provide grade only at a patient or slide level, and are therefore poorly suited to localizing the morphological features that distinguish grades. To enable a patch-level analysis of tumor grade, we use a cohort of 128 slides in which an expert pathologist assigned grades to local tumor regions, complemented by ∼1,000 unannotated whole-slide images that broaden the morphology seen by the generative model (∼1,120 patients total; Methods). This lets us test whether the proposed approach can autonomously nominate discriminative morphological features that recover the same or alternative grading criteria (Fig. 1C).

### Diffusion Autoencoders provide semantically meaningful latent spaces

Following our previous successes of implementing hypothesis-generating IDL for single melanoma cell behavior based on the generative arm of an Adversarial Autoencoder (AAE) model (*60*), we sought to learn similar latent space representations of H&E patches. To adapt the approach to the structurally more complex tissue image patches, we increased the latent space dimension and added an adversarial arm to encourage the reconstructed images to be indistinguishable from the real ones (Fig. 2A). Although the class-agnostically trained AAE produced visually accurate reconstructions (Fig. S2A), the underlying latent space was biologically sub-optimal: image patches with highly similar latent vectors often shared a similar tissue architecture, but could exhibit highly different traits, such as originating from different slides and grades (Fig. 2B, S2B). This phenomenon was true regardless of the latent vector sizes or distance metrics we chose (Fig. S2B and S3A). We reasoned that the discordant nature of the AAE’s latent space arises from the representations having to perfectly recover all aspects of the image, including many trivial aspects such as absolute positions of nuclei that are unrelated to the tumor grade.

**Fig. 2.**
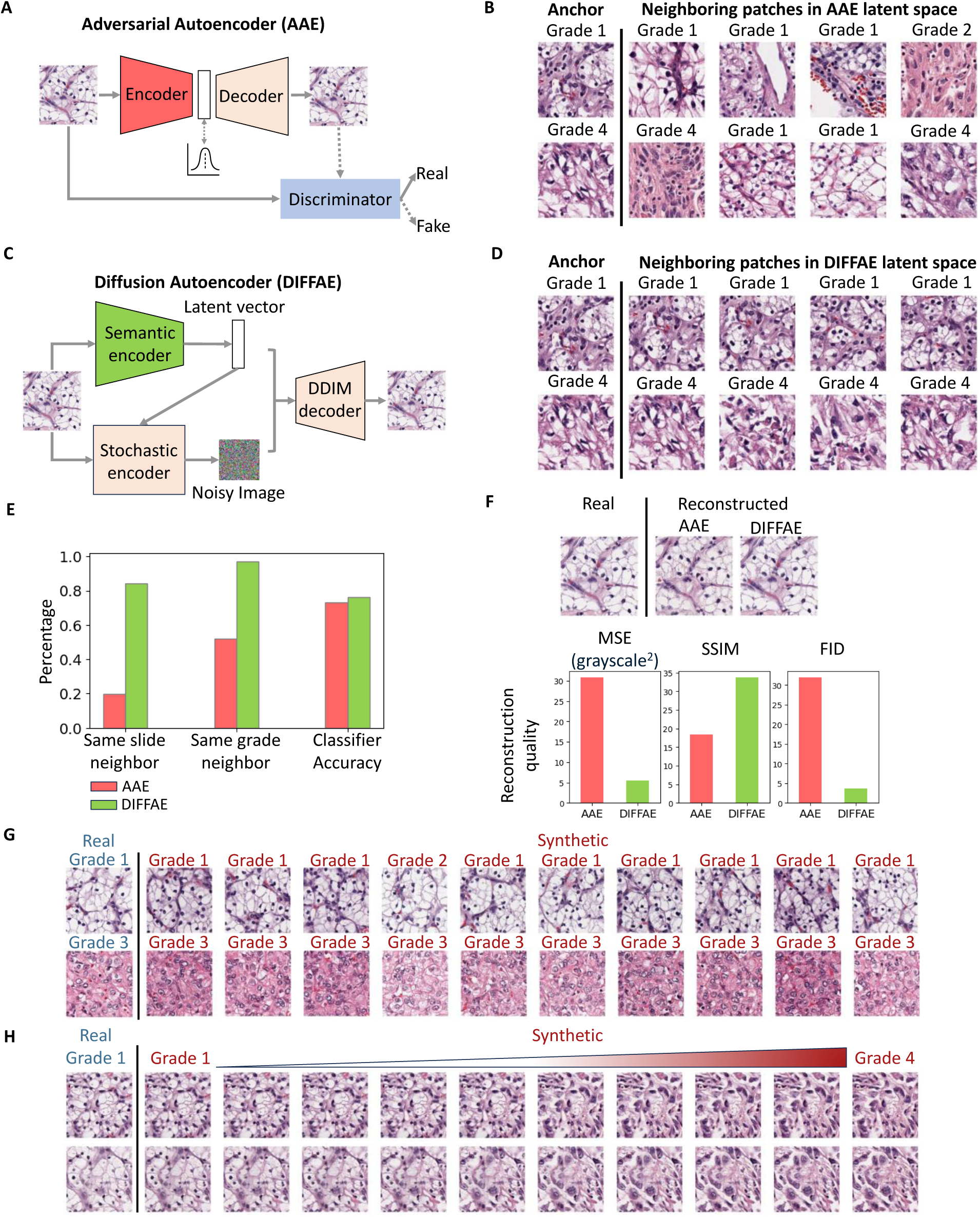
Unsupervised Autoencoder models for the Delta-Marches framework. **(A)** Structure of Adversarial Autoencoder (AAE), which encodes tissues images into a 1024-dimensional latent space and uses a discriminator model to encourage realistic reconstruction. **(B)** Two examples (rows) of patches and their nearest neighbors in the AAE latent space exhibit differing morphology and tumor grades, suggesting the discontinuity in these variables. **(C)** Structure of Diffusion Autoencoder (DIFFAE) comprising a 512-dimensional semantic encoder and a stochastic encoder. **(D)** Examples of neighboring patches in DIFFAE latent space exhibit consistent morphological and tumor grade characteristics, indicating improved latent-space continuity. **(E) C**omparison of the percentage of 10 nearest patches identified using cosine similarity (N=120,000) of a patch which come from the same slides or have the same grade based on the AAE (pink) and DIFFAE (green) latent spaces. LDA accuracy for classifying all four tumor grades in the respective latent spaces. **(F)** Comparison of images reconstructed by AAE and DIFFAE (N=1000 patches), measured using mean squared error (MSE), structural similarity index measure (SSIM), and Fréchet inception distance (FID). **(G)** Demonstration of the interaction between semantic latent vector and noise image of the stochastic encoder. Two tissue images (rows) were encoded into semantic vectors, combined with 10 random noise inputs to synthesize images. Synthetic images on same row share the input tissue phenotype and (with the exception of 1 patch) have the same grade as predicted by VGG19 tumor grade classifier but differ in spatial structure. Images in the same column share the spatial patterns generated by the same noise inputs. **(H)** Demonstration of tumor grade transformation by semantic latent-space traversal while preserving spatial structure. Grade 1 semantic vectors were shifted along the grade 4 classifier axis and combined with unchanged stochastic noise from real images to generate synthetic sequences showing smooth grade-associated morphological transitions.

To overcome the need for specifying low level detail in our latent vector, we adopted diffusion models that are widely used for generating high quality images based on broad descriptive prompts (e.g., DALL-E(*74*)). Diffusion models (*75, 76, 78, 79*) learn to generate the full spectrum of training images by iteratively denoising a random signal, and the prompt vector associated with each image merely serves as a bias directing this process. Specifically, we chose to replace the AAE with a Diffusion Autoencoder (*75*) where the image representation is effectively bifurcated into two parts (Fig. 2C): the latent vector captures the high-level, semantic image content, whereas the low level interchangeable image aspects (such as the absolute locations of nuclei) are delegated to the stochastic encoder. Compared to the AAE, DIFFAE i) generates a more biologically meaningful latent space in which tissue images with similar latent vectors originate from the same slide and grade (Fig. 2D, E); ii) provides better separability of grades based on Linear Discriminant Analysis (LDA) in the semantic latent space (Fig. 2E); and ultimately iii) also improves the quality of image reconstructions, as assessed by visual inspection and standard image quality metrics (Fig. 2F, S2C, S3A). To situate DIFFAE among current approaches, we benchmarked it on two axes: latent-space consistency against the UNI2-h (*86*) encoder, and image synthesis against the generators PixCell (*87*) (conditioned on UNI2-h) and MuPaD (*88*). DIFFAE produced higher-fidelity images than both generators (lower FID; Fig. S3B), while its latent space was modestly less discriminative than the larger, more heavily pretrained UNI2-h encoder (Fig. S3A). Because the Delta-March is agnostic to the underlying representation as long as it is paired with a generative arm, these results support our use of DIFFAE here while leaving room to incorporate stronger representations in future. Having established a meaningful and generative latent space, we used

DIFFAE to generate different tissue images representing the same grade (Fig. 2G) and to simulate the different appearances of the same tissue as it would progress from Grade 1 to Grade 4 (Fig. 2H). The former image series are generated by varying the noise component for a fixed position in the semantic latent space. The latter image series are generated by traversing the semantic latent space (Methods) from regions encapsulating Grade 1 to regions encapsulating Grade 4, leaving the noise component fixed. Importantly, the latent space is learned in a fully unsupervised manner, without grade labels; grade enters only through the traversal direction, the vector of maximal grade separation from a 4-class linear classifier. Traversal along this direction produced a smooth transition across tumor grades in both latent space (Fig. S4A) and image space (Fig. 2), while not altering cohort identity (Fig. S4B, C), supporting that the learned trajectory captures grade-associated morphology rather than cohort-specific variation.

### Evaluation of synthetic tumor grade data by human pathologists

We sought to gauge the realism of the DIFFAE-synthesized images by having experienced pathologists inspect them, blinded to both the source (‘synthetic’ vs ‘real’) and the tumor grade of each tissue patch image. Specifically, using the DIFFAE we generated 40 synthetic images equally split across all four grades (as determined by our grade model) and merged them with a similarly distributed set of 40 real images. We then presented these 80 images one–by-one in randomized order to four pathologists with experience in ccRCC pathology. They were tasked with specifying for each patch the source (real vs synthetic) and grade (low: 1/2 or high: 3/4). The ability to distinguish real from synthetic images was no better than random (accuracies of 45% to 56%) for both low- and high-grade images (Fig. 3A). In addition, inter-pathologist agreement was generally low, with a mean Cohen’s kappa of 0.15. In contrast, on average pathologists achieved 92% accuracy in identifying the grade on both real and synthetic images (Fig. 3B) with ground truth based on our grade model and similar results using individual pathologists to assign grade (Fig S5A). Albeit not statistically significant, lower accuracies were observed in classifying low-grade images both real and synthetic, with about 11% being mistaken for high-grade. This possibly reflects the challenge of grading small patches: as grade calls are based on the most aggressive cells, a small number of cells with more aggressive appearing phenotypes could lead to a high-grade call in a low-grade region. The tumor grading performance was comparable on real and synthetic data both in terms of agreement with the grade model (Fig. S5B, C) and among pathologists, which showed a mean Cohen’s kappa of 0.75 (Fig. 3B). The divergence between these two agreement levels is itself informative: the same raters who reliably reached consensus on grade could not agree on real versus synthetic, indicating that grade leaves consistent, recognizable signatures the synthetic images preserve, whereas synthetic origin leaves no systematic tell. Overall, we conclude that the DIFFAE model generates realistic tissue images that preserve the characteristics of assigned tumor grades.

**Fig. 3:**
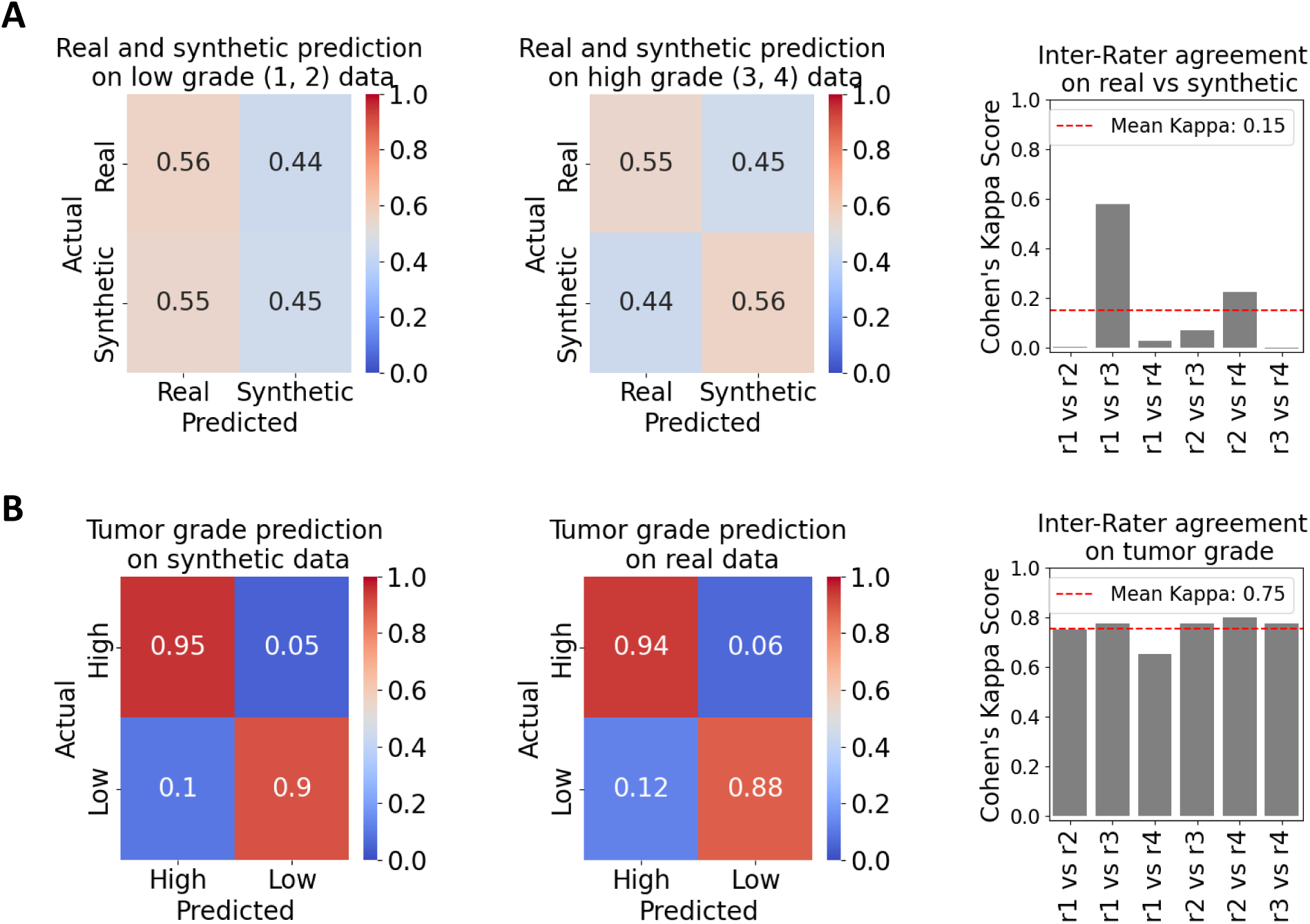
Classification of DIFFAE-generated images by expert pathologists. Four pathologists were tasked with determining for N=80 tissue image patches whether i) the image was real or synthetic and ii) it displays a low grade (grades 1 or 2) or high grade (grades 3 or 4) tumor. **(A)** Accuracy of distinguishing real from synthetic images at low (left) or high (middle) grade. The average accuracy obtained is comparable to a random chance level regardless of tumor grade. Inter-rater agreement across pathologists (r1–r4) is quantified using Cohen’s kappa (right). **(B)** Tumor grading accuracy on synthetic (left) and real (middle) data, along with inter-rater agreement (right). Ground truth grade levels were determined by grade classification model (see Fig. S5 for between pathologist agreement and lowest agreement samples).

### Latent traversals reflect within-patient tumor evolution

Having established the fidelity of individual synthetic images, we next asked whether traversing between two points in latent space generates trajectories that capture biologically plausible evolutionary transformations within a patient’s tumor. The ideal test would compare these synthetic trajectories against real disease progression, but branching evolution and intratumor heterogeneity make ground-truth longitudinal images of solid tumors difficult to obtain. As a proxy, we leverage a multi-region sequencing dataset (WSI1 cohort (*89*)) in which phylogenetic trees have been reconstructed for tumor regions sampled from the same patient. Prior work (*89*) has established that the morphological similarity between such regions largely mirrors their genetic relatedness. This provides a biologically motivated expectation (Fig. 4A): traversal between two regions from the same evolutionary lineage should produce intermediates that resemble other regions on that lineage (positive samples) rather than regions from unrelated branches matched for grade and slide origin (negative samples). We tested this expectation using the pretrained DIFFAE model, noting that neither this cohort nor these patients were part of the DIFFAE training data.

**Fig. 4:**
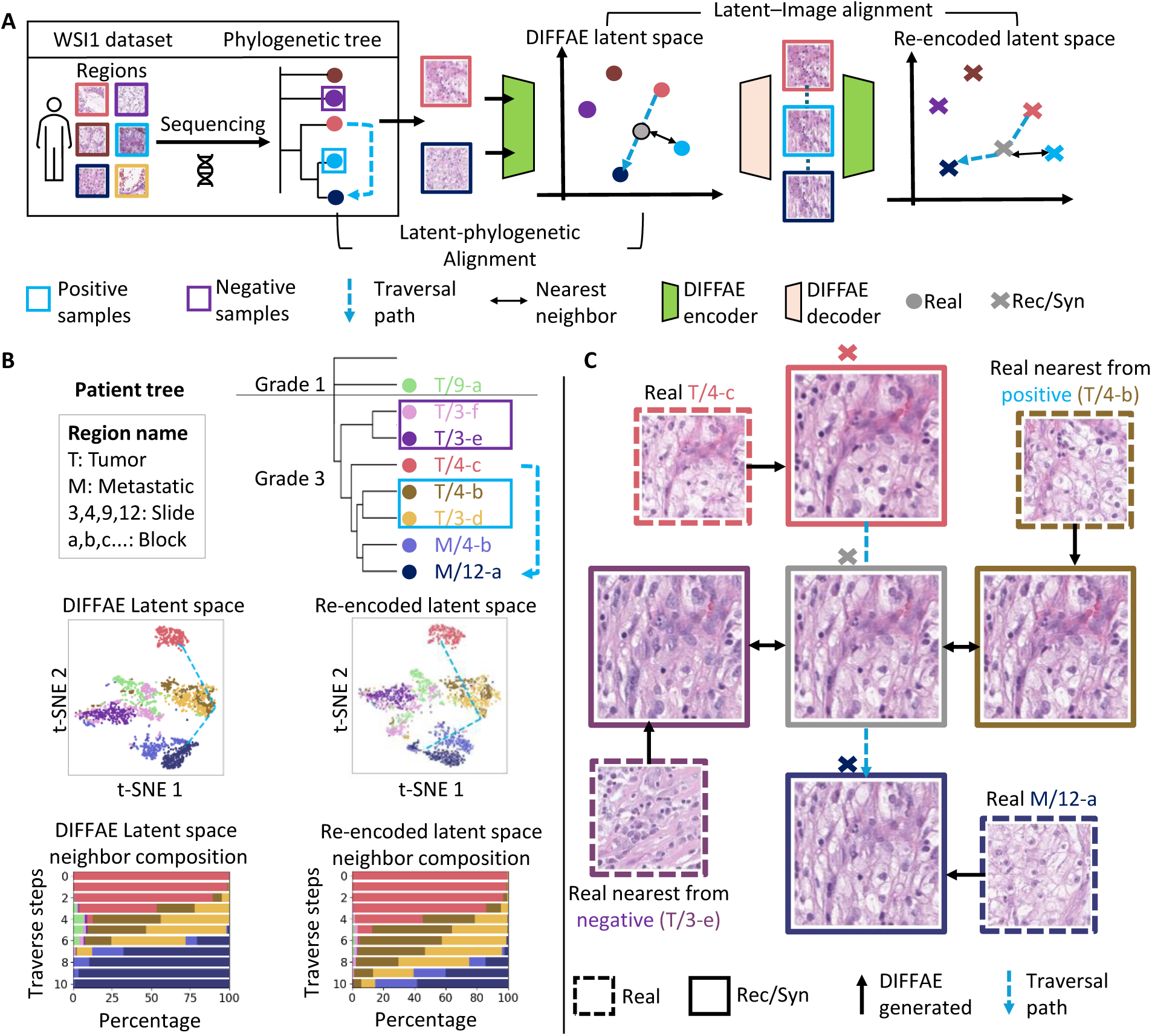
Latent traversal across phylogenetically related tumor regions. **(A)** WSI1 (held out from DIFFAE training) provides matched sequencing data used to reconstruct phylogenetic trees for tumor regions per patient. To evaluate latent-image consistency, real images are first reconstructed by the pretrained DIFFAE model and re-encoded to define a common latent space. Latent traversal (N = 10 steps; dashed arrow) is performed between regions within a major branch. Synthetic images (×) are generated at each interpolation step using the starting image’s DIFFAE stochastic component, then re-encoded into latent space. Similarity is analyzed in two spaces: interpolated versus real-image embeddings in the latent space, and re-encoded synthetic images versus reconstructed-image embeddings in the re-encoded latent space. Positive samples are regions within the same genetic sub-branch, while negative samples correspond to grade- and slide-matched regions from different branches. **(B)** Representative patient analysis. The phylogenetic tree (first row) shows traversal between grade 3 regions T/4-c and M/12-a. Positive samples (T/4-b, T/3-d) are located within the same sub-branch, while negative samples (T/3-f, T/3-e) are selected from different branches with matched grade and slide T/3. Second row: the DIFFAE latent space and the re-encoded latent space of reconstructed images are visualized by t-SNE, with blue dashed lines indicating example interpolation trajectories. For each interpolated and corresponding re-encoded point from a synthetic image, *k* = 10 nearest neighbors are identified in the corresponding latent space using cosine similarity. The neighborhood composition results are summarized in the bar plots. **(C)** Example traversal showing the midpoint synthetic image and its nearest real neighbors from positive and negative regions. For comparison, synthetic images generated from the semantic representations of nearest-neighbor samples, using the same stochastic component, are also shown. Results for additional patients are provided in Fig. S6.

Specifically, we examined patches from different genetic branches of the same patient (Fig. 4B) and compared them to intermediate patches obtained via latent space traversals. To assess traversals, we interpolated between randomly paired patches from two grade-3 regions (T/4-c and M/12-a) rather than along the global classifier direction used for grading, so that both endpoints corresponded to observed samples. Both for individual patches and for the neighbor composition of intermediate points along trajectories, the DIFFAE latent vectors and those re-encoded from the generated images were in close agreement (Fig. 4B, t-SNE and bar plots), indicating that the generated images reflect their intended latent positions rather than drifting off-manifold. This agreement was held even though each traversal paired the stochastic component of one sample with interpolated semantic vectors from another, a more demanding test than simple reconstruction. Although the DIFFAE was optimized for image generation rather than phenotypic similarity, its latent space tracked the underlying biology: regions were arranged consistent with their genetic relationships (Fig. 4B, t-SNE), and trajectory intermediates were consistently enriched for regions within the same evolutionary sub-branch over grade- and slide-matched regions from different branches (Fig. 4B, bar plots). We illustrate this with a representative traversal (Fig. 4C; blue dashed lines in Fig. 4B, t-SNE), in which the generated intermediates exhibit smooth morphological transitions across steps and the nearest real neighbor of the synthetic intermediate belongs to the same genetic lineage, with greater morphological similarity than grade-matched negative controls. Similar results were obtained across additional patients, when intermediates were compared against the latent vectors of real rather than reconstructed images, and under alternative traversal parameterizations (Fig. S6). Together, these results support the biological feasibility of Delta-Marches by showing that DIFFAE-based latent traversals remain on-manifold and consistent with genetically defined tumor relationships.

### Leveraging synthetic image series of grade transitions to nominate grade-driving image features

Our goal for an interpretability-first DL approach is to autonomously identify image features that distinguish specific sample properties of interest – here, cancer grades – and aid pathologists in formulating hypotheses on the underlying property-driving mechanisms. To this end, we developed an analysis referred to as *Delta-Marches* (Fig. 5A) that facilitates the identification of phenotypes linked to changes in the specific sample properties. Rather than comparing different tissue images across different grades, which vary in numerous other aspects besides the grade, Delta-Marches reveal how a particular tissue patch changes solely because of a grade transformation. While one could attempt to quantify the changes between pairs of counterfactual images by subtracting their raw RGB intensities, such differences primarily reflect pixel-level fluctuations in color lacking semantic or structural context. Therefore, the resulting maps highlight nearly all tissue regions—excluding only very dark pixels—thereby overwhelming the image with nonspecific signals and failing to isolate meaningful morphological changes (Fig. S7, S8). Instead, we compare each pair of counterfactual images in a perceptual feature space (*90–92*), measuring how their responses differ across a fixed convolutional filter bank. The specific filters are not critical (quantified below): we use only the first 9 layers of the pretrained AAE (not the full encoder), and the first 13 layers of a supervised grade classifier give similar results (Fig. S7, S8). We subsequently track hotspots of representation changes, captured in a *Delta-Map*, between consecutive steps of the Delta-March.

**Fig. 5.**
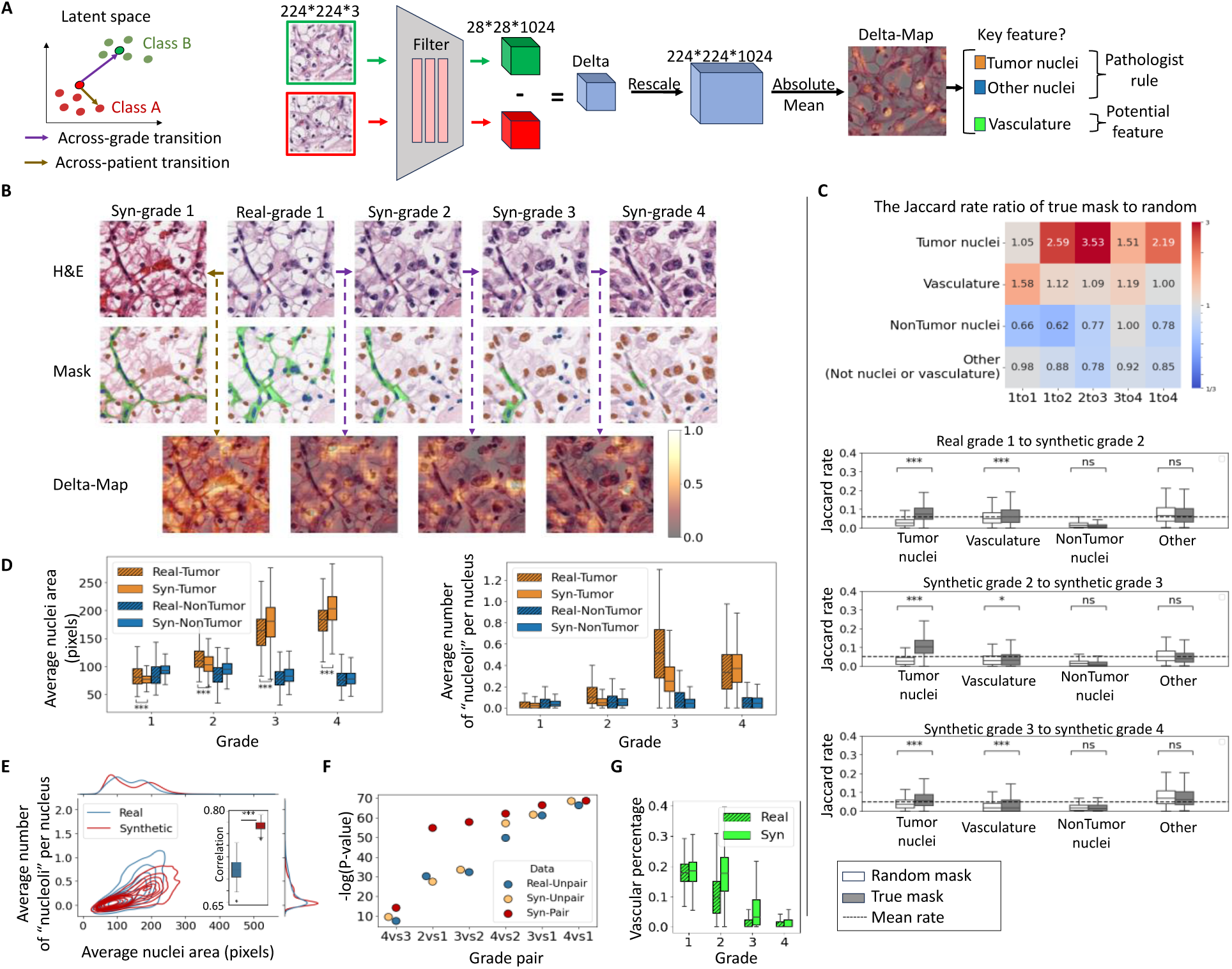
Hypothesis generation on grade-driving mechanisms using Delta-Marches. **(A)** Overview of the Delta-Marches framework to identify spatially distributed and perceptual features that alter during a latent-space traversal across tumor grade. Across-patient transitions serve as negative controls. **(B)** Representative example showing the real/synthetic images along the latent transition (top), corresponding cell-type segmentations (middle), and overlaid Delta-Map intensities between successive steps (bottom). **(C)** Jaccard similarity analysis comparing Delta-Maps to tissue segmentations. For N = 977 grade 1-to-4 transitions, binarized Delta-Maps (threshold = 0.5) were compared with both actual segmentations and random masks using the Jaccard index. The table displays the median ratio of actual-to-random Jaccard indexes. Bar plots show Jaccard indexes for indicated tissue components; dash line is the average across all samples and components in each transition. **(D)** Tumor and non-tumor nuclear phenotypes variation across grades in real and synthetic data (N = 500 patches per grade). Synthetic grades 1–2 come from grade 4-to-1 traversals (N = 556), and grades 3–4 from grade 1-to-4 traversals (N = 977). “Nucleoli” are identified as local extrema in the hematoxylin channel. **(E)** Relationship between average tumor nuclear area and “nucleoli” number in real and synthetic images. Contour plot highlights joint distributions (N = 500), with inset showing correlations from 100 random samples of 100 patches per grade. **(F)** Statistical power to detect grade-associated increase in per-patch average tumor nuclear size. For each grade-pair comparison (N = 100 per grade), patches were randomly sampled from real images (Real-Unpair) or synthetic images from the same (Syn-Pair) or independent (Syn-Unpair) grade 1-to-4 traversals. The sign test compared average nuclear size between groups (N = 100 patches), repeated 1,000 times to compute the mean -log(p-value). **(G)** Analysis of percentage area covered by vasculature across grade (N=500 patches per grade).

The Delta-Maps across grade transitions demonstrate a high degree of spatial modulation with localized hotspots (Fig. 5B). Qualitative comparison with standard class activation methods suggests that this approach resolves fine-grained structures often smeared by CNN-based heatmaps, while consistently highlighting the full distribution of altered phenotypes, avoiding the sparse or fragmented explanations typical of ViT-based approaches (Fig. S1). Intriguingly, visual inspection of these results highlights a focus on tumor nuclei, consistent with grading rules. The significance of these hotspots can be appreciated in comparison to a direct jump from grade 1 to 4 (Fig. S7) and to a within-grade negative control (Fig. 5B left column), where we preserve the noisy image (and hence the position of nuclei) of the initial patch but replace its semantic latent vector with that of a different patch of the same grade.

For a more systematic and unbiased analysis, we cross-referenced the Delta-Maps with segmentations of tumor and other nuclei, as well as the vascular network (*85*) and evaluated the enrichment of the Delta-Maps intensity across these segmented areas. Specifically, we binarized the Delta-Maps into low- and high-intensity regions and calculated the Jaccard similarity index, defined as the ratio of the intersection to the union between the binarized Delta-Map and the segmentation mask. Additionally, we compared this Jaccard index to a random baseline, calculated using segmentation masks from a different randomly selected patch of the same grade. The table in Fig. 5C presents the ratio of Jaccard indexes (actual mask to random mask). Higher ratios indicate stronger alignment of the Delta-Map with the segmented tissue component of interest. The ratios reveal that the Delta-Map produced by marches across grade transitions is substantially elevated at the loci of tumor nuclei, but not at the loci of non-tumor nuclei (e.g. immune cells and other cell types) (Fig. 5C). Hence, the Delta-Marches determines that cancer cell nuclei are a key indicator of grade transitions, as postulated by today’s clinically implemented grading rules (*80–82*). Interestingly, the decreased focus on tumor nuclei between grades 3 and 4 (bar plots Fig. 5C) is consistent with the inclusion of non-nuclear tumor cell phenotypes (e.g., rhabdoid or sarcomatoid features) in calling grade 4 (as opposed to grades 1-3, which are purely nuclear). Negative controls, based on within-grade (across-patient) transitions, show lower and less specific ratios, further demonstrating the unique roles tumor nuclei ought to play in across-grade transitions. Although the Jaccard indexes were relatively high in the other regions, i.e. tissue parts not associated with cell nuclei or vasculature derived from true masks, they were lower than those obtained from the random masks leading to ratios below 1. This suggests that no major features exist in the tissue background that would determine the grade transition.

Although lower than those for tumor nuclei, the Jaccard indexes associated with vasculature across grade transitions were still higher than the indexes from random masks, i.e. the index ratios between true to random masks are consistently above 1 for all grade transitions. This indicates that vasculature may be a secondary marker of tumor grade. Interestingly, small-step marches captured vascular changes, whereas direct large jumps from grade 1 to grade 4 overlooked these subtle features. This underscores the advantages of the Delta-Marches, which integrates changes in tissue appearance over a continuous, controlled progression, allowing the detection of subtle grade-related morphological shifts that may be obscured by tissue heterogeneity and variation agnostic to the grade transition. These conclusions are robust to the choices of perceptual filter and segmentation model and reproduce when the analysis is restricted to either institutional cohort (Fig. S9), indicating they are not driven by batch differences between institutions.

Next, we focused on the putative phenotypic changes in nuclei from across-grade transitions in synthetic tissue image patches (Fig. 5D). We found that tumor cell nuclei, but not nuclei of other cell types, increased in size with grade, highlighting the cell-type awareness of the semantic latent space. Moreover, the count of peaks of hematoxylin intensity per nucleus, which we use as a proxy for prominent “nucleoli” (Methods and Fig. S10), also grew only in tumor cells (Fig. 5D). Notably, the change in nuclear size and nucleoli between grade 3 and grade 4 was relatively less pronounced compared to other grade transitions in both real and synthetic data, reflecting nuclear features are not the primary distinguishing feature for grade 4 classification, consistent with the reduced Jaccard ratio shown in Fig. 5C. This shows that the IDL system based on Delta-Marches reproduces established grading rules directly for the data. Interestingly, the tumor grade effect on nuclei is more pronounced in grade transitions in the semantic latent space, with notably smaller nuclei in synthetic low grades (1–2) and larger nuclei in synthetic high grades (3–4) compared to real data (Fig. 5D). While the individual trends for nuclei and “nucleoli” are largely in agreement between real and synthetic data, the correlation between these two features is stronger in the synthetic data (Fig. 5E, S11). Thus, while real image patches display some of the expected grade related features, these are more consistently present in synthetic data.

We sought to more formally test whether the ability of latent space traversals to solely modulate grade increased the statistical power over traditional cross-grade comparison where other non-grade aspects may also change. Specifically, we wondered to what extent any differences arose from our synthetic trajectories exaggerating grade features as opposed to overcoming interpatient heterogeneity within a grade by following the same patch across grades. To address this, we compared the average nuclear size across tumor grades (1 to 4) using both real and synthetic data. In the real data, images were randomly sampled from each grade group (Real-Unpair). For synthetic data, two approaches were used for each grade-pair comparison: (1) Syn-Unpair, where samples originated from independent latent space transitions, which would allow us to test solely the difference between synthetic and real feature distributions; (2) Syn-Pair, where counterfactual samples were derived from the same latent space trajectories grade 1 to grade 4, thereby potentially overcoming interpatch variability. For each grade pair, we applied the sign test to compare the per-patch average nuclear size between groups (N = 100 patches), repeating the analysis 1,000 times. The modest and inconsistent p-value improvements in Syn-Unpair (Fig. 5F) suggest that feature amplification alone has a limited impact on statistical separation. In contrast, the consistently smaller p-values observed in the Syn-Pair data highlight the power of counterfactual analysis enabled by Delta-Marches to emphasize grade-relevant tissue features more effectively than conventional cross-grade comparisons. Notably, the synthetic pair comparisons produced the most dramatic results relative to those observed in real data for the grade 1-to-2 and 2-to-3 transitions (Fig. 5F), where strong correlations within a trajectory help control heterogeneity. For more distant grades, the correlation between nuclear sizes weakens, and differences are increasingly driven by distributional shifts. While in this case, the nuclear size has a strong enough effect to be detected also in real data, this approach will allow us to pinpoint more complex traits with smaller effect sizes and in more data-limited applications. Taken together these findings highlight two key advantages of the proposed Delta-March: (1) the ability to isolate the effect of individual variables on phenotype changes using analysis of counterfactual images, and (2) the use of small, controlled transitions along a biologically meaningful latent axis to enhance statistical power and provide a more precise and interpretable means of uncovering underlying disease mechanisms.

Finally, we were curious to see whether the reduction in vasculature with grade previously conjectured by us and others (*85*) was also reflected in synthetic image series of latent space transitions in the direction from low to high tumor grades. We found indeed that the model recapitulated this effect to a level comparable to that in real data (Fig. 5G). We note that unlike tumor-nuclei, within grade transitions seemed to impact the vasculature pattern (Fig. 5C, 1st column), possibly because there are non-grade related aspects that also impact the specific vascular pattern observed.

### Validation of Delta-Marches in colorectal tumor progression

To test whether Delta-Marches generalizes beyond ccRCC grading, we applied the framework to colorectal tissue, focusing on the transition from normal mucosa to high-grade dysplasia, a progression characterized by glandular enlargement and distortion, loss of goblet cells, and disruption of normal mucosal organization (*93–95*). We retrained the DIFFAE on the HunCRC colorectal dataset (*96*), which provides annotated patches from multiple classes including normal, low-grade, and high-grade dysplasia which we focus on. Patches in the dataset are 512×512 pixels reflecting a 248×248 micron tissue area, more than double that of our ccRCC application, resulting in a coarser resolution (after down sampling to the 224×224 px resolution used by DIFFAE) that captures glandular architecture rather than subcellular detail. To test robustness, we kept the AAE filter bank for the comparison of counterfactual images unchanged from the ccRCC analysis. Thus, these experiments tested whether Delta-Marches operate across scales and systems. Synthetic trajectories were generated from normal mucosa toward high-grade dysplasia along a latent linear classifier direction, following the same procedure as in ccRCC. The resulting images show smooth morphological transitions across traversal steps (Fig. 6A).

**Fig. 6.**
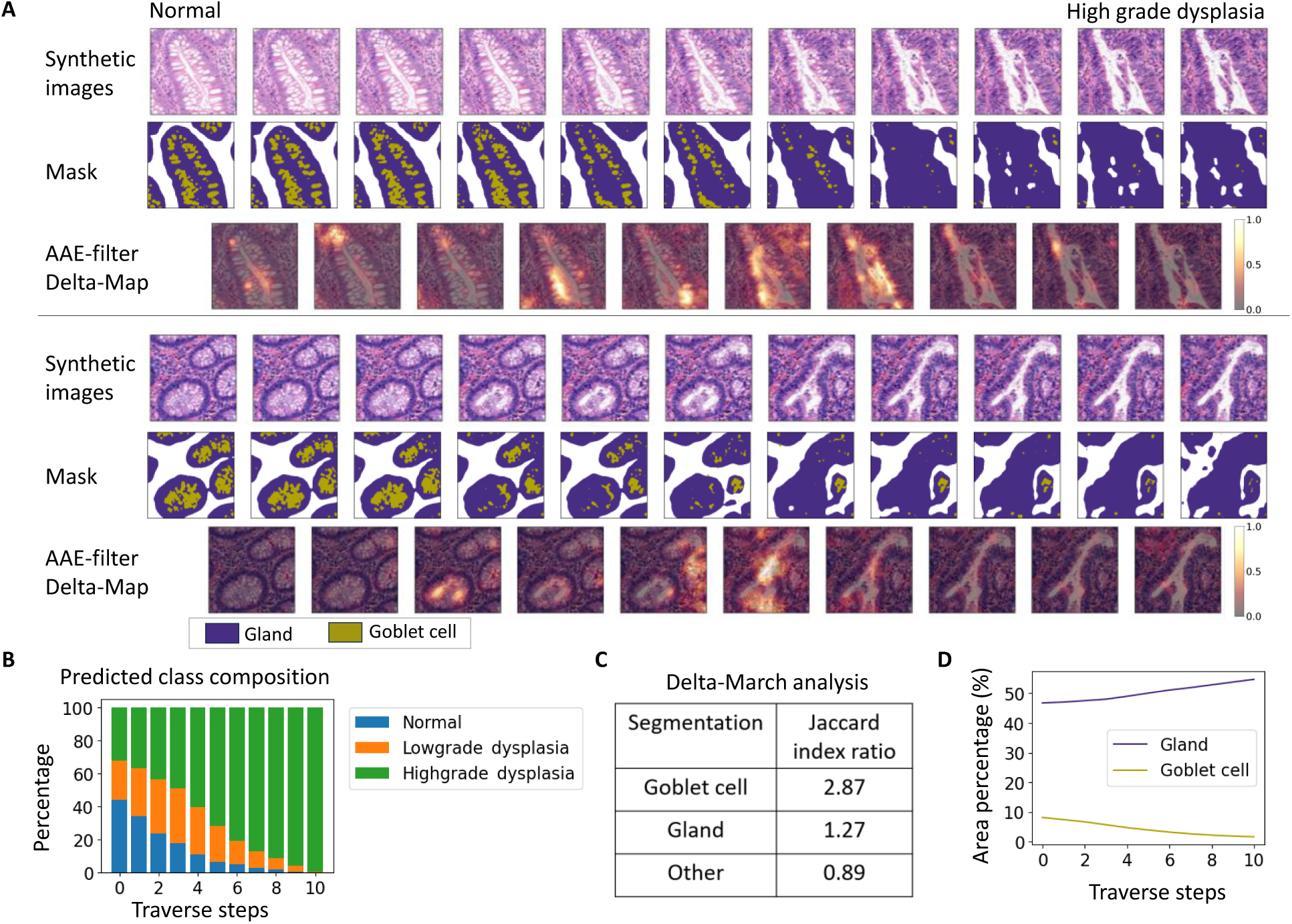
Delta-Marches for colorectal tumor progression. **(A)** Examples of DIFFAE-generated synthetic traversal from normal mucosa toward high-grade dysplasia in colorectal tissue. Rows show synthetic images, gland and goblet cell segmentations, and AAE-filtered Delta-Maps between consecutive traversal points. **(B)** Predicted class composition across traversal steps using a VGG19 classifier trained on real colorectal images from three classes: normal, low-grade dysplasia, and high-grade dysplasia. **(C)** Jaccard index ratio quantifying the overlap between Delta-Maps and segmented gland or goblet cell regions. **(D)** Gland and goblet cell area percentage across traversal steps.

To better understand these transitions, we used the HunCRC data to train a separate model to identify normal/low-grade/high-grade dysplasia. Although the traversal was gated only to begin at normal and terminate at high-grade dysplasia, intermediate steps were classified as low-grade, indicating the latent trajectory recovers the correct ordinal progression rather than interpolating directly between the endpoints (Fig. 6B). Next, we used publicly available, externally developed gland and goblet cell segmentation models (*97, 98*) to localize regions of maximum change. Delta-Marches analysis (Fig. 6C) nominated goblet cell regions as the most prominent sites of transformation (Jaccard index ratio 2.87), with glandular regions also enriched (1.27). Similar to the ccRCC analysis the background tissue showed no enrichment (0.89), suggesting that also in this case there are no major hidden morphological features that would add a notable marker of the normal mucosa to high-grade dysplasia transition. Furthermore, continuous quantification across traversal steps (Fig. 6D) revealed a progressive decline in goblet cell area concurrent with increasing glandular density, recapitulating the morphological correlates of progression from normal mucosa to high-grade dysplasia (*93–95*). These results demonstrate that Delta-Marches recover known progression features in a tissue system with fundamentally different architecture and at a coarser spatial scale than the ccRCC analysis.

## DISCUSSION

There are three fundamental challenges in using real world data to identify meaningful phenotypic associations between tissue morphology and disease properties (e.g. mutation status of a single gene or tumor aggressiveness for which grade is a proxy). First, morphological traits manifest highly non-linearly as image features, and much of the variation in the image pixel space, such as absolute positions of nuclei, is irrelevant. Second, it is rare to find samples differing solely in the disease property of interest, and thus observed morphologic changes across specimens are confounded by differences in non-related properties. Third, even if we could solely change the disease property of interest, large biological changes (e.g. a jump from grade 1 to 4) will impact multifarious aspects of morphology due to the complex inter-connected networks of influence. In attempting to overcome these challenges, we take inspiration from the production of caricatures where cartoonists identify the essence of a subject they wish to highlight and then depict it with a minimal set of strokes. Translating this process into an AI system, we address these challenges by a) using a DIFFAE to produce meaningful latent representations of images consigning irrelevant parts into a noise vector, b) identifying a direction in the latent space such that changes are solely associated with the property of interest, and by c) traversing the latent space in a Delta-March along this direction to identify the most salient affected regions in the image. Thus, the presented framework serves as an AI-powered cartoonist that, when presented with representative data of both consequential and inconsequential variations, autonomously extracts the association between tissue appearance and the property of interest.

Although xAI activation maps and Delta-Marches both highlight areas of importance overlaid on images, they represent fundamentally different concepts and corresponding outputs. Delta-Maps represent generative estimates of counterfactual differences between class-shifted images, an explicit simulation of how tissue morphology would change under an idealized shift in biological state. As a result, Delta-Maps not only carry directionality, but the area highlighted represents the scale of the phenotypes being changed, whether sub-cellular or architectural. In contrast, xAI activation maps measure a pre-defined classifier’s local sensitivity to perturbations in the image. As classifiers are optimized for classification accuracy, the spatial distribution of these maps, whether focused on a single pixel or a broad range, primarily reflects the functioning of the classifier and less so the underlying biological principles. Delta-Marches also differs from recent counterfactual approaches which use generative models to render an image as a different class (*62, 69, 99*), but leave a human to spot the resulting differences and typically flip class in a single jump. Delta-Marches instead quantifies the change as a spatially resolved Delta-Map and integrates it over small steps, surfacing subtle, dispersed features such as the grade-associated vasculature reduction missed by a direct grade 1 to 4 transition (Fig. 5C).

Translating this framework into practice required a representation that isolates the property of interest from irrelevant variation, enabling the generation of image transformations that specifically reflect the intended progression. Our previous generative approach for single cell images (*60*) used a simpler AAE encoder and needed to exaggerate effects beyond the naturally observed variation to highlight pseudopodal extensions as a readout of metastatic propensity. Here in a more complex tissue setting, we used a more sophisticated approach to remove distracting features and highlight features of interest. First, we showed that the DIFFAE’s split of the latent space into meaningful and “noise” aspects (associated e.g., with nuclear positions) results in a more biologically meaningful representation of histopathology images than a simple AAE (*60*). Using the independent multi-region WSI1 dataset, we then showed that latent traversals between a patient’s tumor regions follow that tumor’s genetically defined evolutionary relationships, capturing the biology of an individual tumor rather than average behavior (Fig. 4, S6). Equipped with the confidence from the WSI1 data analysis, we then moved in the latent space of the ccRCC data set to synthesize images of idealized tumor grade transitions while holding other aspects of tumor tissue variation fixed. The resulting synthetic images more consistently exhibited changes to multiple grade-related image features (nuclear size and nucleolar count) than real images (Fig. 5D, E) and even led to a slight increase in pathologists’ grading accuracy on synthetic relative to real images (Fig. 3B). Moreover, comparing nuclei to their synthetically grade warped versions increased statistical power relative to real samples with the same grade difference (Fig. 5F), suggesting the counterfactual modeling enabled by the DIFFAE latent space overcomes sample to sample variation. This could allow detection of subtle effects for which available experimental data does not provide adequate power.

While the proposed strategy is agnostic to data type and biological process, we provide a primary proof of principle demonstration in recovering grading criteria for ccRCC pathology and identifying the key changes in the normal-to-dysplastic colorectal tissue transformation. The Delta-March generates a continuous, controlled model of the increase of grade with single-cell spatial resolution, thereby identifying a minimal set of grade-associated changes. Specifically, in the ccRCC data, the Delta-March process highlighted the tumor nuclei as most impacted, allowing us to then recover the known grading rule of increased nuclear size while the increase in number of local intensity maxima within the nuclear masks mirrors the rule of increased nucleolar prominence. Interestingly, in identifying the decrease of vasculature with grades, which we and others have noted before (*83–85*), our approach goes beyond the formally identified grading rules. VHL loss, an initiating event in ccRCC, causes increased vasculature as cells perceive themselves to be in hypoxic conditions (*100*), but with increasing grade cells can survive without such vasculature, likely due to a metabolic shift away from oxidative metabolism towards the Warburg effect (*101*). Taking together, these results suggest our approach has the capacity to distill the essence of image properties associated with tumor grade change. Beyond ccRCC grading, Delta-Marches pinpointed gland distortion and goblet cell loss as morphological hallmarks of colorectal tissue transition from normal mucosa to high-grade dysplasia. This underscores the potential of the Delta-March concept as a general means of determining decisive morphological characteristics of disease progression.

Future work can build on our work in several ways. As with any method that learns from data, our approach is susceptible to data biases such as a class difference confounded by technical variation (e.g., batch, staining), incomplete representation of classes, or class imbalance, all of which can bias the identification of discriminative features. We find no evidence of this across institutions here, possibly because grade distributions are similarly broad and class balanced across institutions, but future work will need principled means to diagnose and reduce such variation.

Algorithmically, the Delta-March requires only two ingredients: a latent representation that isolates the property of interest, and a generator to render counterfactual images. Here both come from the DIFFAE, but neither is fixed; the representation could instead come from pathology foundation models(*77, 102–104*), and the generator from conditional diffusion models(*76, 105*) such as those transforming images to reflect molecular changes. Similarly, while here we restrict ourselves to interpreting patch level differences, more sophisticated representation strategies will be needed to analyze patient level labels where individual patches may contribute differently. How best to traverse this space is a further open question: linearly as we do or by optimizing some other criterion to surface subtler relationships. More broadly, image classification and generation have traditionally been separate domains. Yet, as the adoption of machine learning algorithms in biomedical domains is limited by their interpretability rather than their classification performance, we anticipate that the development of representations that enable both classification and generative visualization of this space will become important. Delta-Marches would be especially valuable in cancer because of the multifactorial root causes of disease progression and their manifestation in complex morphological signals. Other diseases with similar traits include neurodegeneration, where different strains (protein assemblies differing in structure) have been hypothesized to underlie complex aggregate morphologies that are the gold standard for disease classification (*77, 102–104*). Ultimately, Delta-Marches offers clinicians and researchers a framework to uncover how tissue morphology relates, often in unexpected ways, to disease, advancing both our understanding of disease mechanisms and the development of more personalized treatments.

## MATERIALS AND METHODS

### Data

All datasets used in this study are summarized below; the last two are detailed in their respective sections. All three ccRCC cohorts were analyzed as 224×224-pixel patches extracted at 20X magnification (∼0.5 microns per pixel) from tumor regions of H&E-stained WSIs:

1. Grade cohort: Comprising 128 slides, including 100 slides from UTSW and 28 slides from the Mayo Clinic (*106*) repository, all with localized grade annotation. An expert pathologist (V. Panwar) annotated large tumor regions of coherent morphology and assigned each a grade (1–4), providing a more precise ground truth than the patient-level grade used in the clinic. Patches were randomly sampled within these regions and inherited the grade of their parent region. The per-grade distribution of cases and patches is shown in Table S1.
2. Additional Training Cohort: Comprising additional 1,000 WSIs from the Mayo Clinic repository, as previously described (*106*). Tumor regions were automatically identified using a pretrained region classification model, and patches were randomly sampled within these regions.
3. WSI1 multi-region cohort: a multi-region sequencing cohort (*89*) drawn from three ccRCC patients used to validate that latent traversals reflect within-patient tumor evolution (Results and Methods below).
4. HunCRC colorectal cohort: a publicly available annotated colorectal dataset (*96*) used to test generalization across tissue type and scale (Results and Methods below).

The collection and analysis of the UTSW portion of the Grade cohort and the WSI1 multi-region cohort were approved by our Institutional (UT Southwestern Medical Center and Parkland Hospital, Dallas) Review Board under protocols STU 022015-015 and STU 012011-190 and were conducted in accordance with the Declaration of Helsinki. Archival pathology specimens and retrospective clinical records were analyzed in de-identified form under an IRB-approved waiver of written informed consent because the research involved existing materials and posed minimal risk to participants. Three patients in the WSI1 cohort-Patients A, B, and C-underwent additional genomic analysis. Patients B and C provided written informed consent permitting public sharing of their genomic data; data from Patient A were not publicly released.

### Adversarial Autoencoder (AAE)

#### Architecture

The AAE model adopts an unsupervised framework (*60*), employing a convolutional encoder-decoder architecture. The encoder compresses image through five convolutional blocks, incrementally increasing channel sizes (256, 512, 1024, 1024, and 1024), each followed with batch normalization and Leaky ReLU activation layers. Latent space representations are generated via a fully connected layer. The decoder reverses this process, reconstructing the original image via transposed convolutional layers, batch normalization, and sigmoid activations. Following the original AAE paper (*107*), our model uses an adversarial component intended to make the latent distribution indistinguishable from a normal distribution. Additionally, to enhance the reconstruction quality of high-resolution tissue images, we added a discriminator, similar to those used in Generative Adversarial Networks (GANs), that ensures the real and synthetic images were indistinguishable. This discriminator has five convolutional layers with batch normalization and Leaky ReLU activations, concluding with a fully connected layer and sigmoid output.

#### Training

Training was unsupervised, as is standard for autoencoders. However, to ensure that the model sees a wide distribution of grades, for training and testing, we utilized 120,000 patches, with 30,000 patches per tumor grade derived from the Grade cohort. Training involved minimizing a composite loss function comprising reconstruction, adversarial, and discriminator loss terms, balanced for optimal performance. The model was trained for 30 epochs with a fixed learning rate *r* = 10^−4^ for the AAE and *r* = 10^−5^ for the discriminator, respectively.

### Diffusion Autoencoder (DIFFAE)

#### Architecture

We adopted the DIFFAE model (*75*). Briefly, the architecture of a diffusion autoencoder integrates a semantic encoder, a diffusion process, and a decoder to create a robust generative framework. The semantic encoder compresses input data into a latent representation, which is then progressively corrupted through a series of controlled noise-injecting steps in the forward diffusion process. The decoder reverses these steps, refining the latent representation to reconstruct the original input. For our experiments, we retained all default parameter values from the original implementation, with adjustments made solely for the input image dimensions (224×224 pixels).

#### Training

Two distinct training protocols were utilized. For the comparative analysis (Fig. 2) against the AAE model, we trained the DIFFAE model on a subset of 8,000 patches subsampled from the 120,000 patches used in AAE training, evenly distributed across tumor grades. For the final model used in downstream analyses, we trained the unsupervised DIFFAE model on an extended dataset of 18,000 images: 8,000 annotated patches from the Grade cohort and 10,000 unannotated tissue patches extracted from the Additional Training Cohort to capture broader tissue structural variability.

#### Image grade classification and CAM

We trained a grade classifier with a VGG19 backbone on 120,000 patches over 50 epochs with a learning rate of 10^−4^, achieving 92% accuracy on the Grade Cohort training data. The VGG19 model was pretrained on the ImageNet dataset and fine-tuned for four-grade classification using cross-entropy loss.

Class activation map (CAM) is a widely used technique for generating heatmaps that visualize important regions influencing deep learning model predictions. To validate the performance of CAM-based approaches on detecting critical features related to tumor grade, we performed GradCam++, FullGrad, and AblationCAM for our pretrained VGG19 grade classifier focusing on the last convolution layer. These CAM-based models were implemented using the pytorch_grad_cam library.

We additionally sought to test if these results extended to non-CNN architectures. To this end, we trained a Vision Transformer (ViT) on the same dataset, which reached slightly higher accuracy (∼97%, also on the training data). We visualized the model’s focus using ViT-Explain, based on the attention rollout process (*59, 108*) (Fig. S1).

#### DIFFAE latent space transitions (ccRCC Grade)

Our methodology for class transitions is based on the attribute manipulation technique from the DIFFAE framework(*75*). Specifically, in addition to the unsupervised autoencoder, DIFFAE framework(*75*) introduces a linear classifier trained on latent vectors from the semantic encoder to identify the axes that separates classes. Attribute manipulation is then performed by traversing the semantic vector along the corresponding axis. In our work for tumor grade in ccRCC, we applied this approach to 8,000 annotated patches across four tumor grades, analyzing the projection of latent spaces onto classifier axes (Fig. S4). Based on the classifier projections, to transition towards higher grades, we adjust the latent representation linearly along the direction of the grade 4 weight vector, while to transition towards lower grades, we shift it along the grade 1 weight vector.

To simulate progression from grade 1 to higher grades, we manipulate the semantic latent vector while keeping the stochastic component fixed, as obtained from the semantic and stochastic encoders of DIFFAE. The stochastic component (noisy image) is obtained from the real grade 1 image by applying the forward diffusion process up to *T* = 250, a mid-range diffusion depth that introduces substantial Gaussian corruption in fine-scale details while retaining coarse spatial structure. By fixing this stochastic component, we ensure that the generative process is anchored to the original tissue architecture; denoising with a shifted semantic vector thus yields a “warped” version of the original image that retains spatial correspondence while exhibiting the target grade phenotype. The semantic vector is then standardized and incrementally shifted along the grade 4 weight vector in step sizes proportional to the square root of the vector’s dimensionality. This step size, used in DIFFAE(*75*) for isotropic random walks in 512-dimensional Euclidean space, ensures smooth progression between grades while maintaining the meaningful structure of the tissue representation. The shifted vector is subsequently de-standardized and passed to the DIFFAE decoder along with the original stochastic noisy image to generate synthetic tissue images. The transition halts either when the generated image is classified as grade 4 by the pretrained VGG19 tumor-grade classifier or after a maximum of four steps. Upon determining the endpoint, 20 intermediate semantic nodes are interpolated along the transition path. These nodes, along with the original noisy image, are input to the DIFFAE decoder to generate synthetic images depicting the across-grade transition. The tumor grades of these generated images are validated using the VGG19 classifier.

#### Synthetic ccRCC grade-transition datasets

For the ccRCC grade transition analysis, generating a synthetic image from a source of the same grade is not equivalent to generating it from a different grade, as the former constitutes simple reconstruction while the latter requires active transformation. To ensure all synthetic images were produced through the same generative process, we exclusively generated target phenotypes by traversing real samples starting from the opposite grade extreme. A similar strategy was used for colorectal tissue progression, but not for WSI1 multi-region dataset, where transitions were defined between phylogenetically related regions rather than class extremes (details are shown in the sections below). Specifically, we generated synthetic tumor-grade images for two purposes:

1. Pathologist evaluation (Fig. 3): We applied this strategy to real samples from all four grades. Transitions pushed toward grade 1 and grade 4 were used to produce the requisite low and high-grade representations, respectively (Table S2).
2. Identification of grade associated features (Fig. 5): To generate high grade samples, we traversed 5,000 real grade 1 samples, confirmed by both pathologist and model prediction, toward grade 4. Of these, 977 trajectories transitioned across all four grades (1–4) and the highest-confidence images were selected for each grade. Conversely, for low grade samples we reversed the process by shifting 5,000 real grade 4 patches toward grade 1, yielding 556 synthetic samples spanning all grades. For within-grade transitions (across patients), we generated synthetic pairs where the stochastic encoding of a tissue sample was preserved, but its semantic vector was replaced with that of a random tissue sample of the same grade from a different patient.

Additional visualizations of latent transitions are shown in Fig. S7 and S8.

#### Pathologist evaluation on ccRCC tumor grade datasets

Four pathologists with experience in ccRCC were provided with written instructions before independently reviewing images through a web-based form during a 30-minute evaluation session. Images were presented one at a time, and pathologists were blinded to image source. For each image, they answered two binary questions: (1) whether the image was real or synthetic, and (2) whether the image represented low-grade tumor, defined as grade 1–2, or high-grade tumor, defined as grade 3–4.

The evaluation was performed in two batches of 40 images each, for a total of 80 images. Images were balanced across real and synthetic sources and across tumor grades. Pathologists were not shown examples of real or synthetic images before the evaluation and were not instructed to use predefined criteria for either task.

For the real-versus-synthetic task, ground-truth labels were known by construction. For the low-versus-high grade task, the reference labels used in Fig. 3 and Fig. S5B, C were determined by the VGG19 grade classification model. As a sensitivity analysis, inter-rater agreement among the four pathologists was quantified using Cohen’s kappa for both tasks (Fig. 3). We summarized agreement levels across images based on the percentage of pathologists making the same call (100%, 75%, 50%, 25%, or 0%; Fig. S5C) and repeated the grading accuracy analysis using pathologist-assigned grades as alternative reference labels (Fig. S5A). In the latter analysis, each pathologist’s grade call was used in turn as the reference, with accuracy computed from the remaining pathologists’ calls. This yielded accuracies of 89% for synthetic images and 88% for real images, compared with 93% and 91%, respectively, using VGG19-defined labels.

#### Deep Learning filter for Delta-March

To identify phenotypes linked to class changes, we developed the Delta-Marches analysis pipeline (Fig. 5A). This pipeline uses a “perceptual” (*90–92*) CNN filter derived from the first 9 layers of our pretrained AAE. Each image pair is passed through this filter, generating 28×28 feature arrays with 1024 channels. These feature arrays are subtracted to compute Delta arrays, which are then upscaled to match the original image dimensions of 224×224. The absolute values of the resulting arrays are averaged across all 1024 channels to produce Delta maps, highlighting areas of morphological difference between the two input images. Additionally, we also evaluated the Delta-Maps derived from the first 13 layers of our pretrained VGG19-based tumor grade classifier following a similar pipeline (Fig. S7, S8). We selected this depth for Delta-Marches filter to capture feature maps that retain low-level structural details while beginning to incorporate high-level tumor-grade features. To demonstrate the necessity of a CNN filter for capturing morphological feature changes, we computed a naïve baseline that subtracts raw RGB pixel intensities and reports the mean absolute difference across channels (Fig. S7, S8).

#### Delta-Marches transitions

We initially focused on the 977 across-grade transitions (described above) from grade 1 to grade 4, which pass through intermediate grades 2 and 3. To capture fine-grained morphological changes, the transitions were subdivided into sub-transitions: real grade 1 to synthetic grade 2, synthetic grade 2 to synthetic grade 3, and synthetic grade 3 to synthetic grade 4. We also extended the Delta-Marches analysis to across-patient transitions by swapping semantic vectors of tissue samples between different patients of the same grade, enabling a deeper understanding of tumor grading features.

#### Jaccard localization analysis (ccRCC)

To identify key tissue components influencing tumor grading we calculate the enrichment in intensity of the Delta-Maps across tumor nuclei, non-tumor nuclei, vasculature, and other (not nucleus or vasculature). Nuclei classification and vasculature segmentation masks for a single grade image were derived using corresponding models developed by the Rajaram lab (*85*). The nuclei segmentation pipeline classifies pixels into three classes: 2 = tumor nuclei, 1 = non-tumor nuclei, and 0 = non-nuclei. The vasculature segmentation model assigns pixels as either 1 = vasculature or 0 = non-vasculature. Composite, across-grade masks were derived as follows:

1. Merging: Nuclear/Vascular masks from the initial and final tissue images were merged by assigning each pixel the maximum label present across both images.
2. Artifact Removal: We identified “ghost nuclei” (nuclei appearing in one mask but not the other, typical of generative artifacts) and eliminated them from the combined mask to prevent biological misinterpretation. The frequencies of the “ghost nuclei” are shown in Fig. S12.
3. Integration: The merged nuclei and vasculature masks were integrated into a single four-class map: 3 = tumor nuclei, 2 = non-tumor nuclei, 1 = vasculature, and 0 = background. To prevent bias in overlapping regions, pixels classified as both vasculature and nucleus were randomly reassigned to a single class.

To quantify the localization of morphological changes to specific cell types, we calculated the Jaccard similarity index (intersection over union, IoU) between binarized Delta-maps and the segmentation masks as follows. First, the Delta-map values were linearly normalized to the [0,1] range per patch and values greater than 0.5 were set to 1 and the rest to 0. The Jaccard similarity index for each component was computed as: 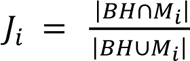, where *M_i_*, *i* = 0,1,2,3 represents the binary segmentation of component *i* (3 = tumor nuclei, 2 = non-tumor nuclei, 1 = vasculature, 0 = other), and BH denotes the binary Delta-Map in each sample. Here, “other” refers to all pixels outside tumor nuclei, non-tumor nuclei, and vasculature, including stroma, whitespace, and unannotated background.

To establish a baseline for comparison, we computed the Jaccard index against segmentation masks from randomly selected patches within the same sub-transition grade. Next, we computed ratio of true and random Jaccard indexes for each patch and reported the median value across all patches for a grade transition (Table in Fig. 5C). Additionally, we assessed the statistical significance of enrichment in specific cell types by comparing the Delta-map similarity against the random baseline, using a one-sided T-test (Bar plot in Fig. 5C).

For comparison with traditional interpretable deep learning approaches, we repeated the same analysis using activation maps from Grad-CAM++ (VGG19 classifier) and ViT-Explain (ViT classifier) on 4,000 real images (1,000 per grade). Results are shown in Fig. S9.

As an independent robustness analysis, we repeated the Delta-Marches analysis using HoVer-Net-derived nuclei masks (*109*), with nuclei labels assigned by majority voting from the Rajaram lab pixel-level nuclei classification mask. Results are shown in Fig. S9C.

#### Impact of institution (ccRCC)

The Grade cohort combines slides from two institutions (UTSW and Mayo Clinic), raising the possibility that grade-associated features could be confounded by institution-associated technical variation. We therefore assessed institutional effects at both the latent-space and Delta-Marches levels.

At the latent-space level, we examined DIFFAE features from the Grade cohort stratified by institution. We encoded Mayo Clinic patches (N = 1,847) and UTSW patches (N = 6,153), visualized their DIFFAE-extracted features using t-SNE, and compared the kernel density distributions of cosine projections onto the grade 4 latent traversal vector across institutions and tumor grades (Fig. S4B,C). While the t-SNE showed visible institutional separation, the grade 4 direction ordered tumor grades consistently within each institution.

At the Delta-Marches level, we repeated the Jaccard analysis separately within the Mayo and UTSW patches of the Grade cohort, drawing the random baseline from the same institution and otherwise following the procedure described in the Jaccard localization section above. The enrichment of Delta-Map signal at tumor nuclei, including its peak at the grade 2-to-3 transition and its decline at grade 3-to-4, was reproduced in both subsets (Fig. S9D), indicating that the grade-associated localization is not driven by a single institution.

#### Nucleoli detection (ccRCC)

Since standardized nucleolus detection algorithms are unavailable, we developed a heuristic approach that identifies “nucleoli” as intensity peaks in Hematoxylin-stained sections of H&E images. We first convert the image to the HED color space and isolate the hematoxylin channel, normalizing and smoothing it using a Gaussian filter with standard deviation *δ* = 1 to enhance nucleoli detection. We then identify nuclei on the nuclei mask, distinguishing between non-tumor and tumor nuclei based on our pretrained nuclei segmentation. “Nucleoli” are detected as local maxima (prominence threshold *α* = 0.08) in the smoothed hematoxylin image, representing darker, separate spots within the nuclei. However, as nuclei themselves represent local peaks in hematoxylin intensity, this approach initially detected “nucleoli” in nearly all nuclei (Fig. S10). To correct this, we subtract a count of one from each nucleus with detected nucleoli. Finally, we calculate the average number of “nucleoli” per nucleus for each type, providing quantitative nucleoli statistics.

#### Counterfactual analysis on real and synthetic data

To highlight Delta-March’s ability to improve statistical power in detecting underlying biological mechanisms, we performed a counterfactual analysis (Fig. 5F) examining average nuclear area across tumor grades 1 to 4. We analyzed latent traversal from grade 1 to grade 4 with 977 corresponding patches per grade. For real data, we sampled 977 patches per grade to maintain comparability with synthetic data. Samples for each grade-pair (N = 100 patches per group) are obtained through three strategies: (1) Real-Unpair: Randomly sampled image patches from each grade group using the real dataset, (2) Syn-Pair: Synthetic image patches for each grade pair derived from the same random latent trajectory, thus preserving progression continuity in the synthetic generation, (3) Syn-Unpair: Synthetic image patches from independent latent trajectories, simulating samples from unrelated transitions. To quantify differences in nuclear areas, we applied the sign test to compare the per-patch average nuclear area between the two grades. For Syn-Pair, samples were paired by trajectory; for Real-Unpair and Syn-Unpair, samples were randomly paired one-to-one between grade groups before applying the same test. The entire process was repeated 1,000 times to assess the robustness and consistency of the statistical signal, measured by the mean log-transformed P-value.

### Latent traversal on WSI1 dataset

#### Data

WSI1 is a morphology-guided multi-regional sequencing dataset ccRCC(*89*) comprising 26 morphologically coherent regions from three patients. For each region, H&E imaging was paired with genetic sequencing, which was used to reconstruct each tumor’s phylogenetic relationships, providing a genetic reference for within-patient evolution against which the latent traversals are compared. Multiregional sequencing was performed across 18 WSIs, with regions distributed as follows: Patient A, 8 regions (6 primary [T] and 2 small-intestine metastatic [M] sites); Patient B, 7 regions (5 primary [T] and 2 pancreatic metastatic [M] sites); and Patient C, 11 regions (8 primary [T] and 3 metastatic [M] sites). In the region names, the numeric identifier denotes the slide and the lowercase letter denotes the tissue block within that slide. From each region we randomly sampled 250 image patches (224×224 pixels at 20× magnification), yielding 6,500 patches in total.

#### Pipeline

Latent traversal on the WSI1 dataset was performed using two approaches. In the sample-across approach (Fig. 4, S6), we randomly constructed 250 image pairs from the traversal regions and performed interpolation between the semantic latent vectors of each pair using 10 interpolation steps. In the region-across approach (Fig. S6), we used the vector between the centroids of two regions, with 250 sampled patches per region. For both approaches, the stochastic component was obtained from the real image at the origin of the traversal path using the DIFFAE stochastic encoder. For the region-across approach, the final semantic point was defined as:

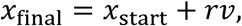

where *x*_start_ is the semantic vector of the real origin image, *v* is the centroid vector between the start and end regions, and *r* = 1.2 is a scaling factor. We used *r* = 1.2 to extend the traversal path slightly beyond the direct centroid-to-centroid distance, allowing the final points to better reach the target region distribution (Fig. S6).

To assess whether traversal intermediates resemble phylogenetically related regions, we performed a nearest-neighbor analysis in the DIFFAE latent space using cosine similarity. For each interpolated latent point, the k = 10 nearest real-image embeddings were identified in the original latent space. For each re-encoded synthetic image, the k = 10 nearest reconstructed-image embeddings were identified in the re-encoded latent space (Fig. 4, S6), and the k = 10 nearest real-image embeddings were also identified in the original latent space (Fig. S6). At each traversal step, neighborhood composition was summarized as the fraction of nearest neighbors assigned to each tumor region considered for the corresponding patient. We then compared this composition to the regions’ phylogenetic relationships, using the phylogenetic trees reported in the original publication (*89*), to test whether intermediates were preferentially closest to regions on the same evolutionary lineage (Fig. 4, S6).

### Delta-Marches on colorectal dataset

#### Data

For the colorectal analysis, we used images from the published HunCRC dataset (*96*), with training data characteristics summarized in Table S3. The unsupervised DIFFAE model was retrained on this dataset. Original patches were extracted at 512 × 512 pixels at 20× magnification and resized to 224 × 224 pixels for DIFFAE training.

#### Pipeline

Latent traversal was performed using the same attribute-manipulation strategy as in the ccRCC tumor grade analysis. Briefly, a linear classifier was trained on DIFFAE semantic latent vectors using the colorectal classes listed in Table S3. The resulting classifier axis corresponding to high-grade dysplasia was used to generate trajectories from normal mucosa toward high-grade dysplasia. We initialized traversal from 1,000 normal samples and generated 10 interpolation steps for each trajectory. The stochastic component was obtained from normal samples by the stochastic encoder. As in the tumor grade analysis, a VGG19 classifier was trained on real colorectal images from three classes—normal, low-grade dysplasia, and high-grade dysplasia—and was used to classify synthetic images and define stopping criteria. The latent transition stopped once the image was classified as high-grade dysplasia, or after two traversal steps, to ensure realistic images.

To characterize morphological changes along the traversal, we applied published gland and goblet cell segmentation models. The gland segmentation model (*98*) was designed for 512 × 512 patches at 20× magnification, whereas the goblet cell model (*97*) was developed for 640 × 640 images at 10× magnification. Accordingly, DIFFAE-generated images were resized to 512 × 512 for gland segmentation. For goblet cell segmentation, the synthetic images were whitespace-padded to 640 × 640. To assess whether the Delta-Marches filter generalized across tissue contexts, the AAE-based filter trained in the ccRCC analysis was kept unchanged and was not retrained on the colorectal dataset. The Jaccard rate ratio in Fig. 6 was computed at each traversal step (N = 10) using the same procedure as in the tumor-grade analysis, and values were averaged across N = 1,000 traversal samples.

## SUPPLEMENTARY MATERIALS

Figure S1-S12, Table S1-S3.

## Acknowledgments

We thank the patients who provided tissues that enabled this research project and are grateful to the Kidney Cancer Program for their support and assistance. We acknowledge BioHPC at UT Southwestern Medical Center for providing the computational infrastructure used in this work. We also acknowledge the insightful feedback and suggestions provided by the Rajaram Lab and the Danuser Lab, especially Dr. Felix Zhou for pointing out the potential usefulness of the DIFFAE model.

## Author Contributions

Conceptualization: T.N., G.D., and S.R.

Computational analysis: T.N., G.D., and S.R.

Model development: T.N., V.J., and A.P.

Data curation: V.P., V.J., and P.K.;

Result evaluation: V.P., C.D., Q.C., and P.K.;

Writing – review & editing: T.N., PK, G.D., and S.R.

Supervision: G.D. and S.R.;

Funding acquisition: G.D. and S.R

## Data and Materials Availability

All data needed to evaluate the conclusions in the paper are present in the paper and/or the Supplementary Materials. The Mayo Clinic WSIs used here (Additional Training Cohort and the Mayo portion of the Grade cohort) are available at https://doi.org/10.25452/figshare.plus.19310870. The colorectal (HunCRC) dataset was obtained from https://doi.org/10.6084/m9.figshare.c.5927795.

The WSI1 dataset was obtained from prior work (89) in the Rajaram lab, where the phylogenetic relationships used here as the genetic reference are also reported. As described there, whole-exome sequencing for the consenting patients (Patients B and C) is available in the European Genome-phenome Archive (EGA) under study accession EGAS50000001064, while the corresponding images cannot be deposited publicly because not all patients consented to unrestricted sharing. Similarly, the UTSW portion of the Grade cohort cannot be publicly released. We are committed to transparency and will make every effort to provide deidentified image patches from the WSI1 and UTSW Grade cohorts upon reasonable request, contingent on appropriate institutional approvals, execution of any required material transfer agreements, and compliance with IRB guidelines. The WSI1 dataset and UTSW portion of the Grade cohort can be provided by Payal Kapur pending scientific review and a completed material transfer agreement through the UT Southwestern Medical Center. Requests should be submitted to.

The DIFFAE model was trained on a compute node equipped with 4× NVIDIA A100 GPUs, requiring approximately 21 hours for 18,000 patches. Inference is substantially less computationally intensive; latent encoding and traversal image generation can be performed on a single GPU, such as an NVIDIA V100. Upon acceptance, all code and trained models, including a Jupyter notebook to generate all figures (attached as supplementary material for review), will be archived in a permanent public repository (Zenodo), with the Rajaram Lab GitHub repository provided as an additional resource.

