## Supplements for "*Delta-Marches:* Generative AI based image synthesis to decode disease-driving morphologic transformations"

### Supplementary Materials for

#### *Delta-Marches*: Generative AI based image transformation to learn histopathology rules

Thuong Nguyen *et al.*

**This PDF file includes:**

Figs. S1 to S12
Tables S1 to S3

**Fig. S1.**

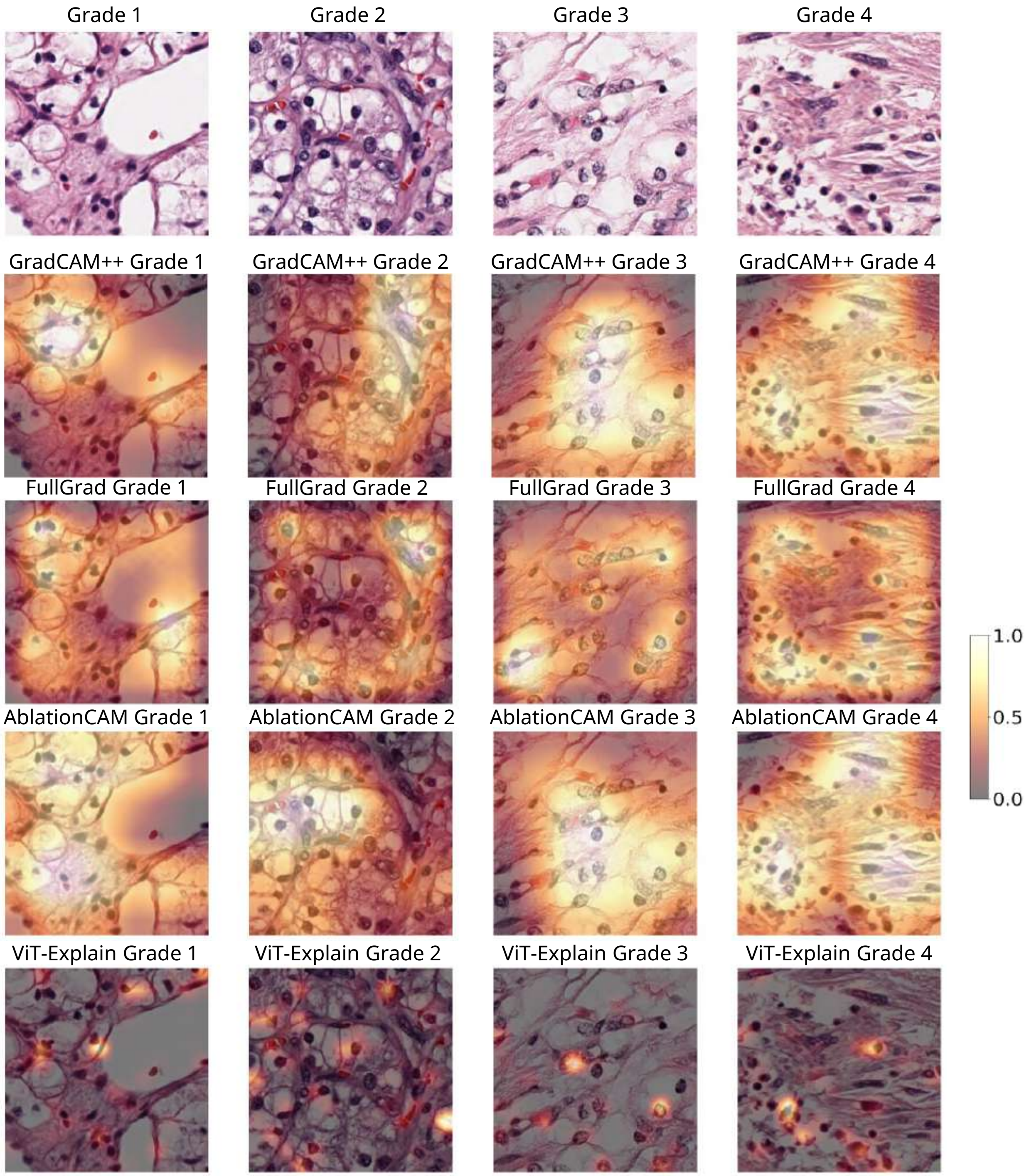


**Fig. S1. Existing interpretability methods recover grade-relevant regions only as broad, nonspecific heatmaps (CNN class-activation maps) or as overly sparse, focal highlights (ViT attention), neither of which isolates the local, spatially dispersed features relevant to tumor grading.** The first row shows the original tissue patches. Rows 2–4 display heatmaps generated from our pretrained VGG19 classifier using GradCAM++, FullGrad, and AblationCAM applied to the last convolutional layer. The final row shows ViT attention maps obtained by attention rollout (ViT-Explain), which instead spotlight a minimal set of highly discriminative tokens, leaving biologically identical tumor nuclei unhighlighted.

**Fig. S2.**

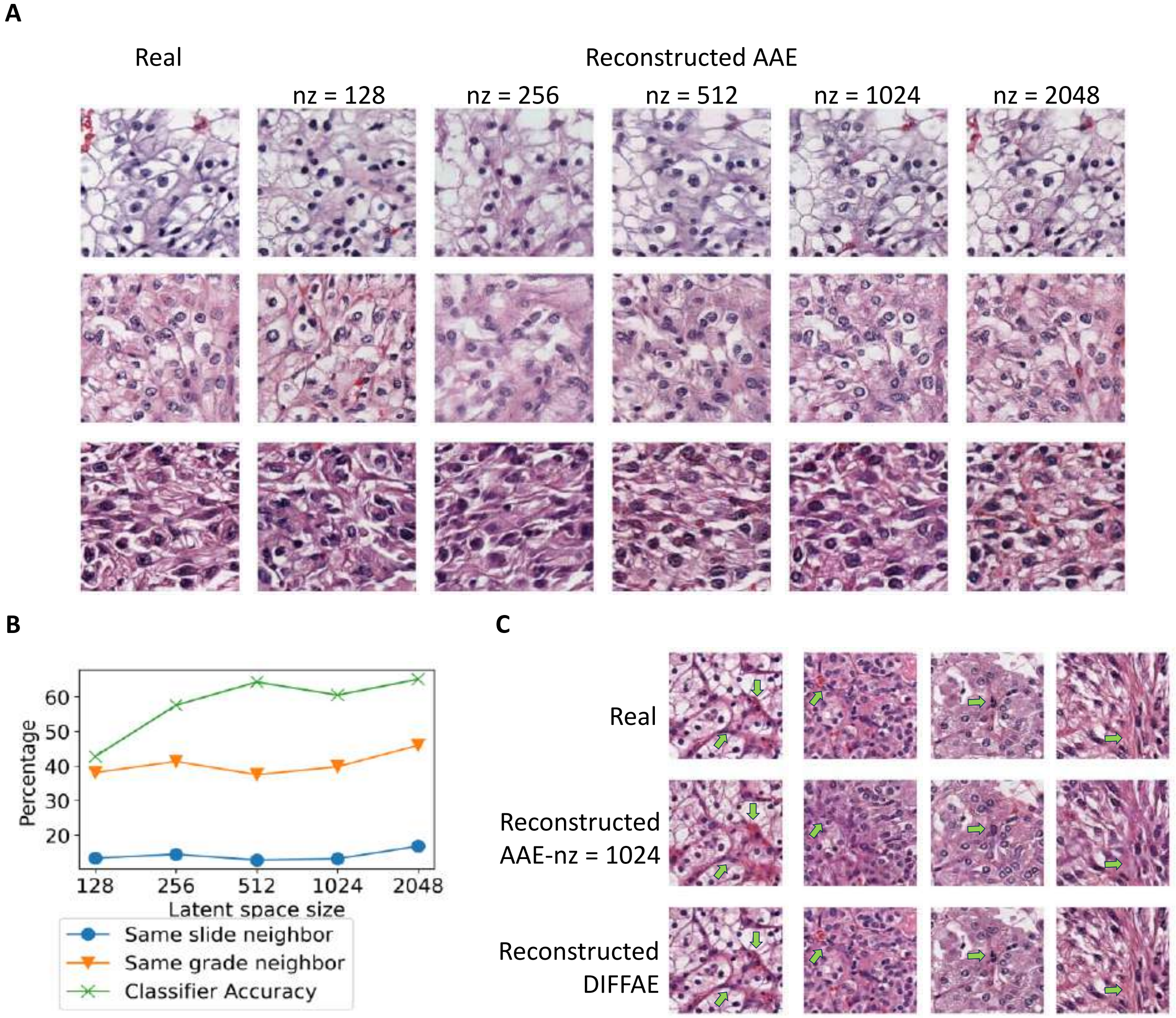


**Fig. S2. Performance of the Adversarial Autoencoder (AAE) in reconstructing histopathological images.** **(A)** Examples of 3 images (rows; leftmost column denotes real images) reconstructed using the AAE model with varying latent space dimensions (columns, nz = 128, 256, 512, 1024, 2048. **(B)** Effect of AAE latent space dimensionality on latent space quality as assessed by different metrics. Blue/Orange plots: average percentages of neighboring patches of a patch having the same slide/grade respectively. Green plots: LDA classifier accuracy in each latent space (N=120,000 patches) **(C)** Additional examples (beyond Fig. 2F) of reconstructed images obtained by AAE model with 1024-dimensional latent space and DIFFAE model. While quality is high in both autoencoders, there are several subtle but biologically significant differences highlighted by arrows.

**Fig. S3.**

A

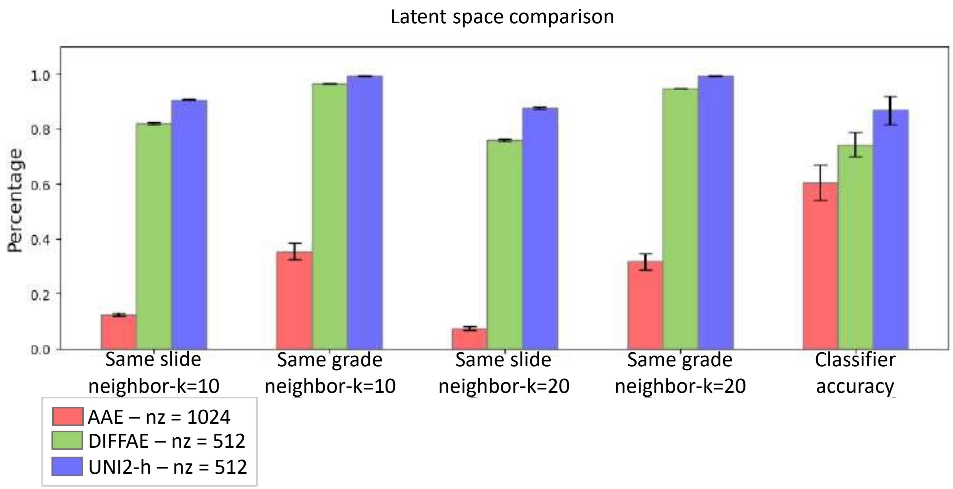


B

| Model | FID |
|---|---|
| DIFFAE | 26.97 |
| PixCell-256 | 38.54 |
| MuPaD-256 | 34.29 |

C

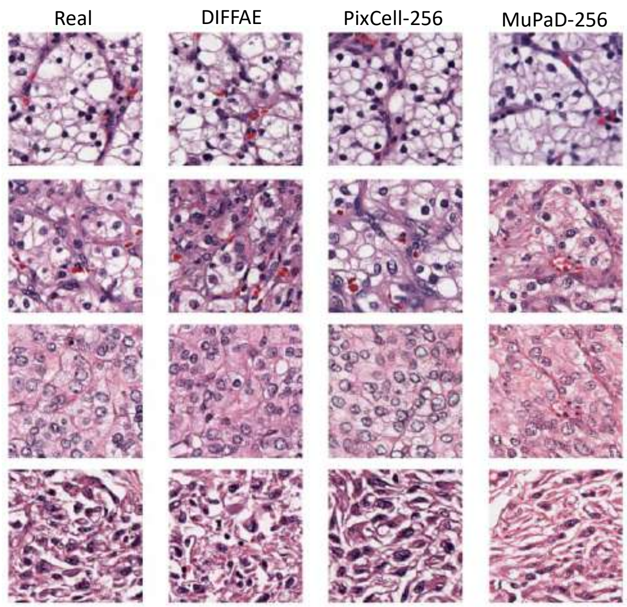


**Fig. S3: Comparison of representation and generative performance of DIFFAE to other approaches. (A)** Comparison of local neighborhood consistency in latent space. For each patch (N = 120,000), the top k nearest neighbors (k = 10 or 20) were identified in the AAE, DIFFAE, and UNI2-h latent spaces using three distance metrics: Manhattan, cosine, and Euclidean distance. Bars show the mean proportion of neighbors sharing the same slide or grade label, and error bars indicate variance across distance metrics. Discriminative performance was further assessed using LDA classifier to separate 4 tumor grades, with accuracy reported as the 5-fold cross-validation average, with error bars indicating variance across folds. **(B)** FID comparing synthetic images from DIFFAE, PixCell-256, and MuPaD-256 against N = 4,000 real images. All images were centercropped and resized to 224×224. FID was computed using a pretrained Inception v3 network. **(C)** Representative real and generated samples from each model.

**Fig. S4.**

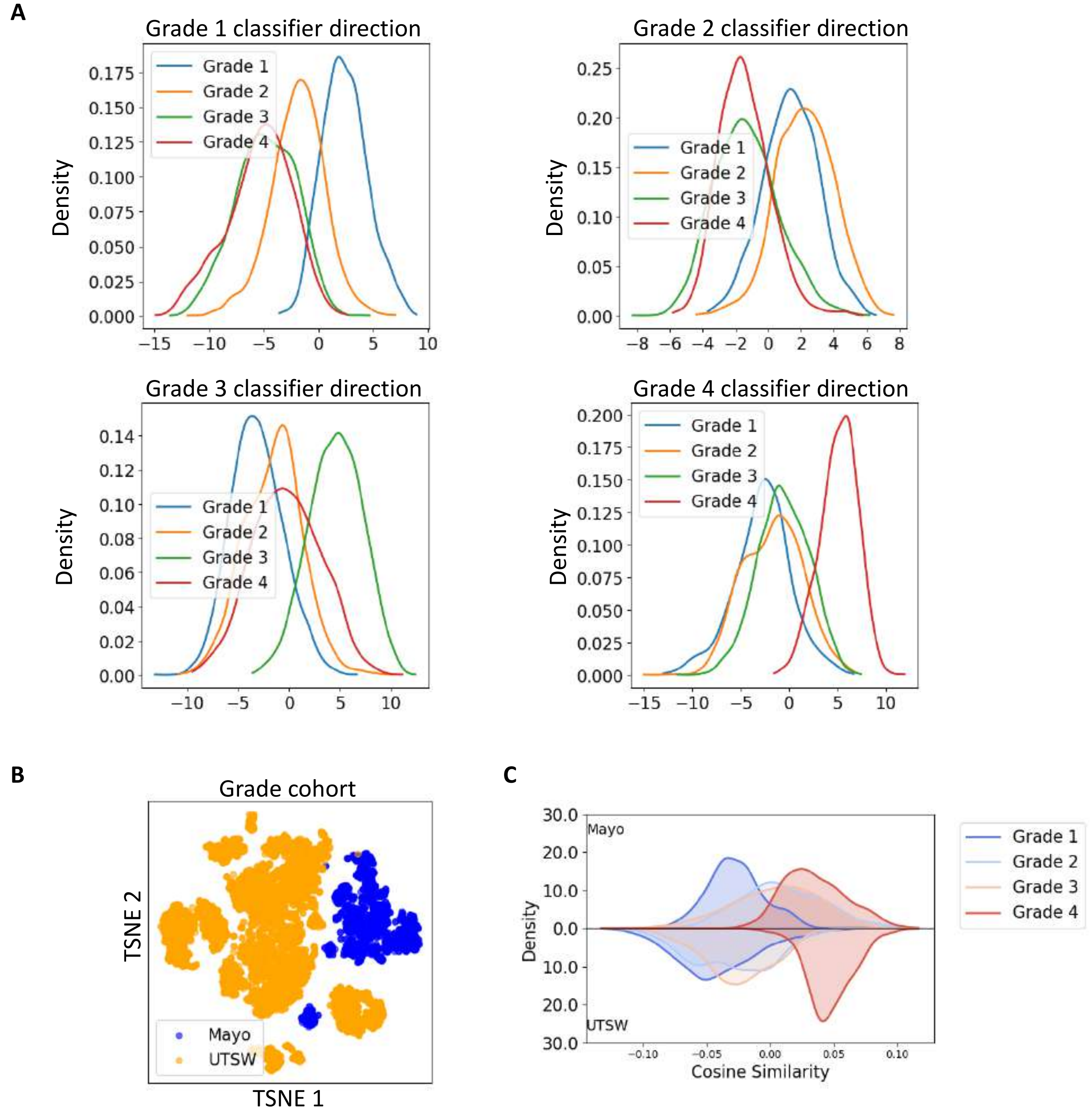


**Fig. S4. Projection of the DIFFAE latent space onto classifier weight vectors, and institution-associated variation in the latent space. (A**) Kernel density estimation (KDE) distributions are obtained by projecting the latent space projections from the training Grade cohorts (N=8,000), onto the weight vectors of a linear classifier. The classifier was trained on DIFFAE semantic latent vectors extracted from the Grade cohort (N=8,000). Each panel illustrates the discriminant score distributions for grades 1 through 4. The grade 4 vector shows a continuous progression from grade 1 to grade 4, while the grade 1 vector displays an inverse trend. **(B)** t-SNE visualization of DIFFAE-extracted features from the Grade cohort, colored by institution: Mayo Clinic (N = 1,847 patches) and UTSW (N = 6,153 patches). The two institutions occupy largely distinct regions, indicating that the latent space captures institution-associated technical variation alongside biological signal. **(C)** KDE distribution of cosine projections of DIFFAE semantic vectors from the Grade cohort (N = 8,000) onto the grade 4 latent traversal vector, stratified by institution (Mayo, top; UTWS, bottom) and tumor grade. Within both institutions the grade 1–4 distributions shift consistently along this direction, showing that the grade axis is shared across institutions and is not driven by the institutional variation seen in (B).

**Fig. S5.**

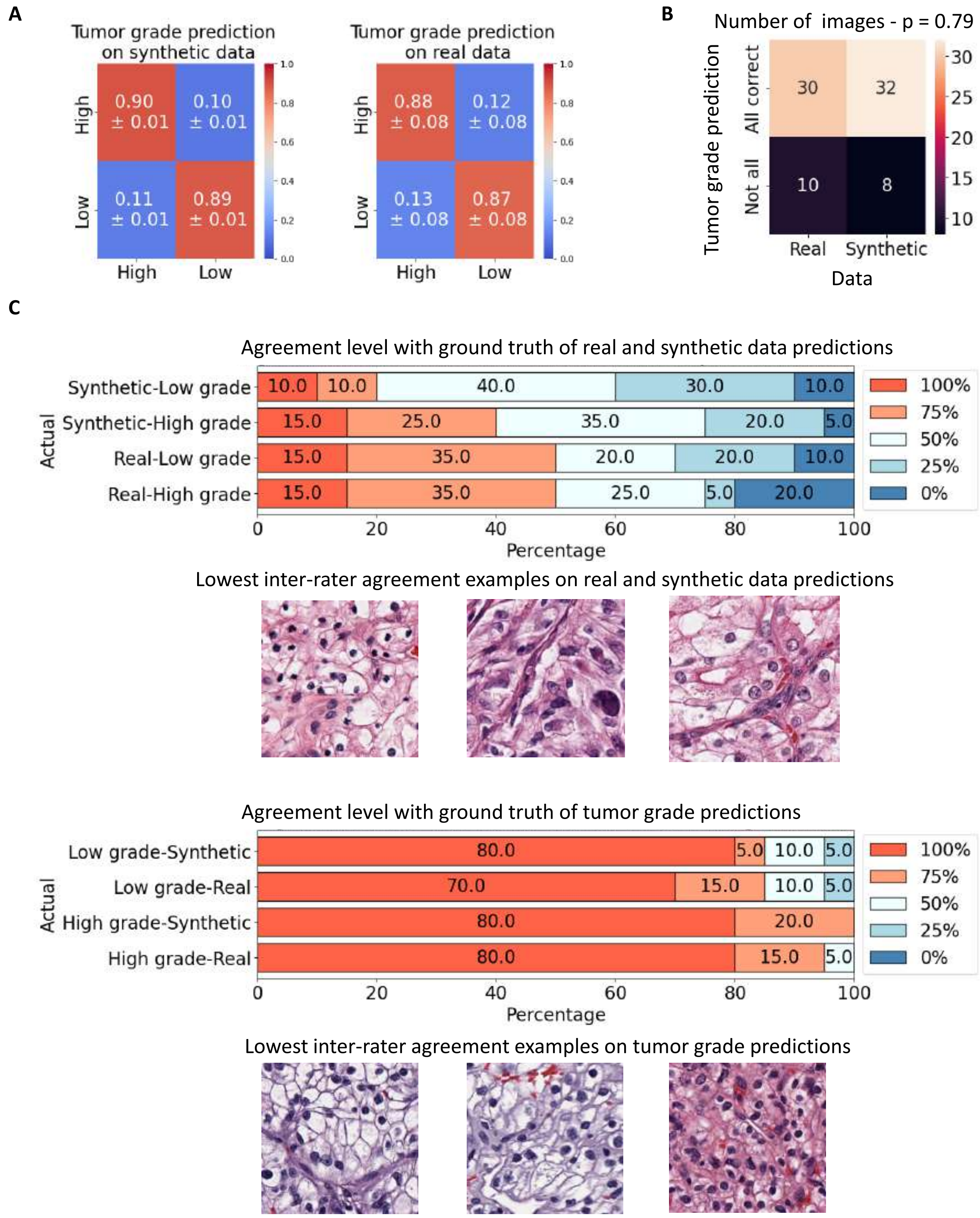


**Fig. S5. Synthetic data captures grading nuances, enhancing pathologists' tumor grade prediction.** We conducted a survey using 80 images, comprising 40 real pathological images and 40 synthetic images generated by DIFFAE models. Each group (real and synthetic) includes 10

patches per grade. Pathologists were tasked with assessing whether the image was (i) real or synthetic and (ii) low (Grade 1, 2) or high (Grade 3, 4) grade. **(A)** Tumor grade accuracy on synthetic (left) and real (right) data, with ground truth based on pathologist-assigned grades (as opposed to the grade model which was used in Fig. 3B). Mean and standard deviation are calculated across ground truth based on individual pathologist with scoring by the others **(B)** Matrix depicting the distribution of correct tumor grade calls by all pathologists in real vs synthetic data (N=80). P-value (shown in the title) is computed using Fisher's exact test to assess differences between the results of the two data groups. **(C)** For each ground-truth group (rows), stacked bars show the percentage of images correctly called by all four, three, two, one, or none of the pathologists (100%, 75%, 50%, 25%, 0% agreement with ground truth), for the real-vs-synthetic call (top) and the high-vs-low grade call (bottom). Grade ground truth is from the classification model. Images below each panel are the maximally split (50%) cases.

**Fig. S6**

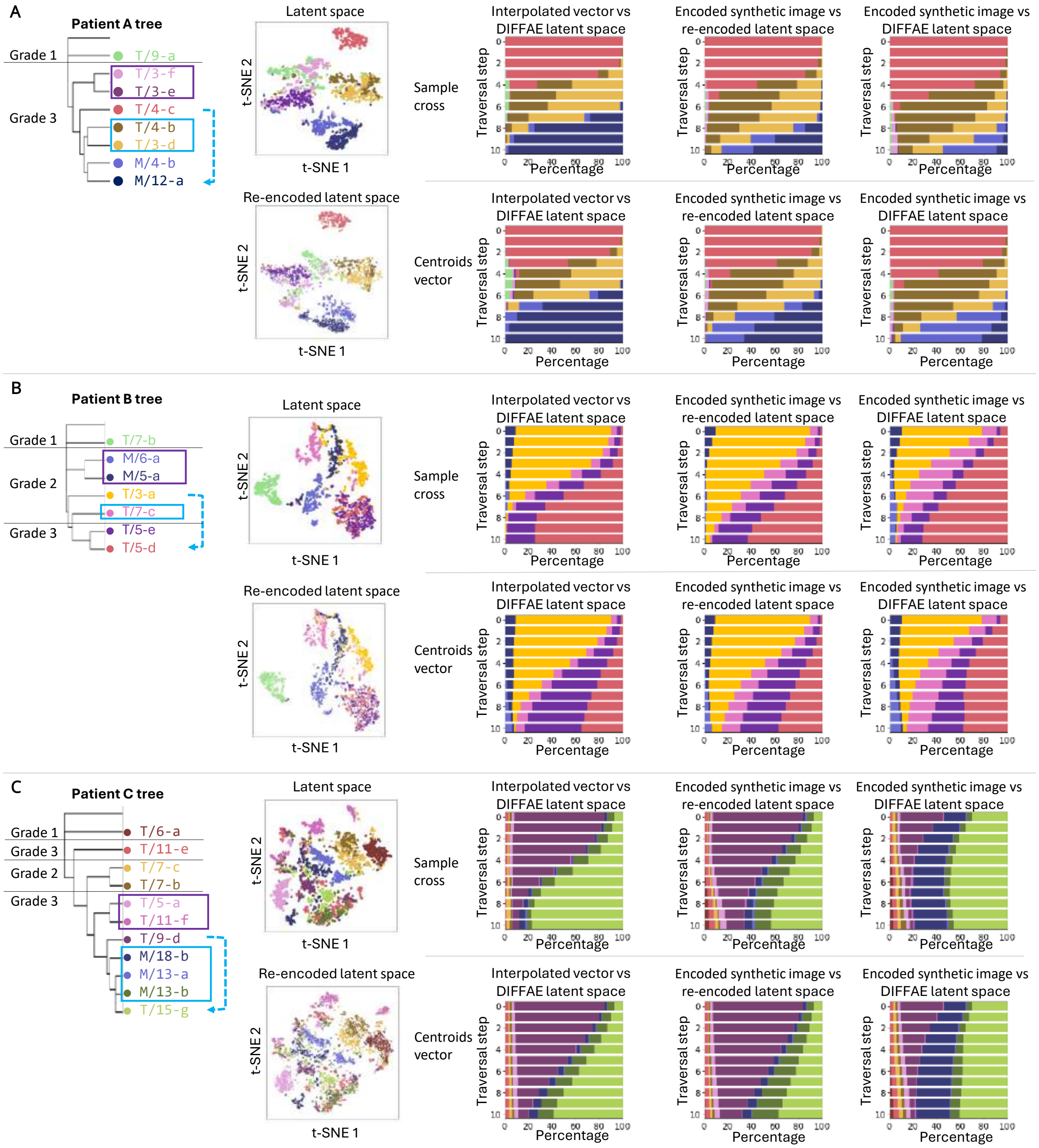


**Fig. S6. Additional examples (beyond Fig. 4) from representative patients.** For each patient, the phylogenetic tree is shown alongside the latent space organization visualized by t-SNE. Latent traversal is performed between tumor regions (N = 250 images per region). Positive samples (blue box) are selected from the same sub-branch, while negative samples (purple box) are drawn from different branches with matched grade and slide. For each point along the traversal and its corresponding synthetic sample, the $k$ = 10 nearest neighbors are identified in the DIFFAE and re-encoded latent spaces using cosine similarity, and the neighborhood composition is summarized in the accompanying bar plots. Traversals were performed using two strategies: across individual samples (N = 250) or using the vector defined by the centroids of the two traversal regions.

**Fig. S7.**

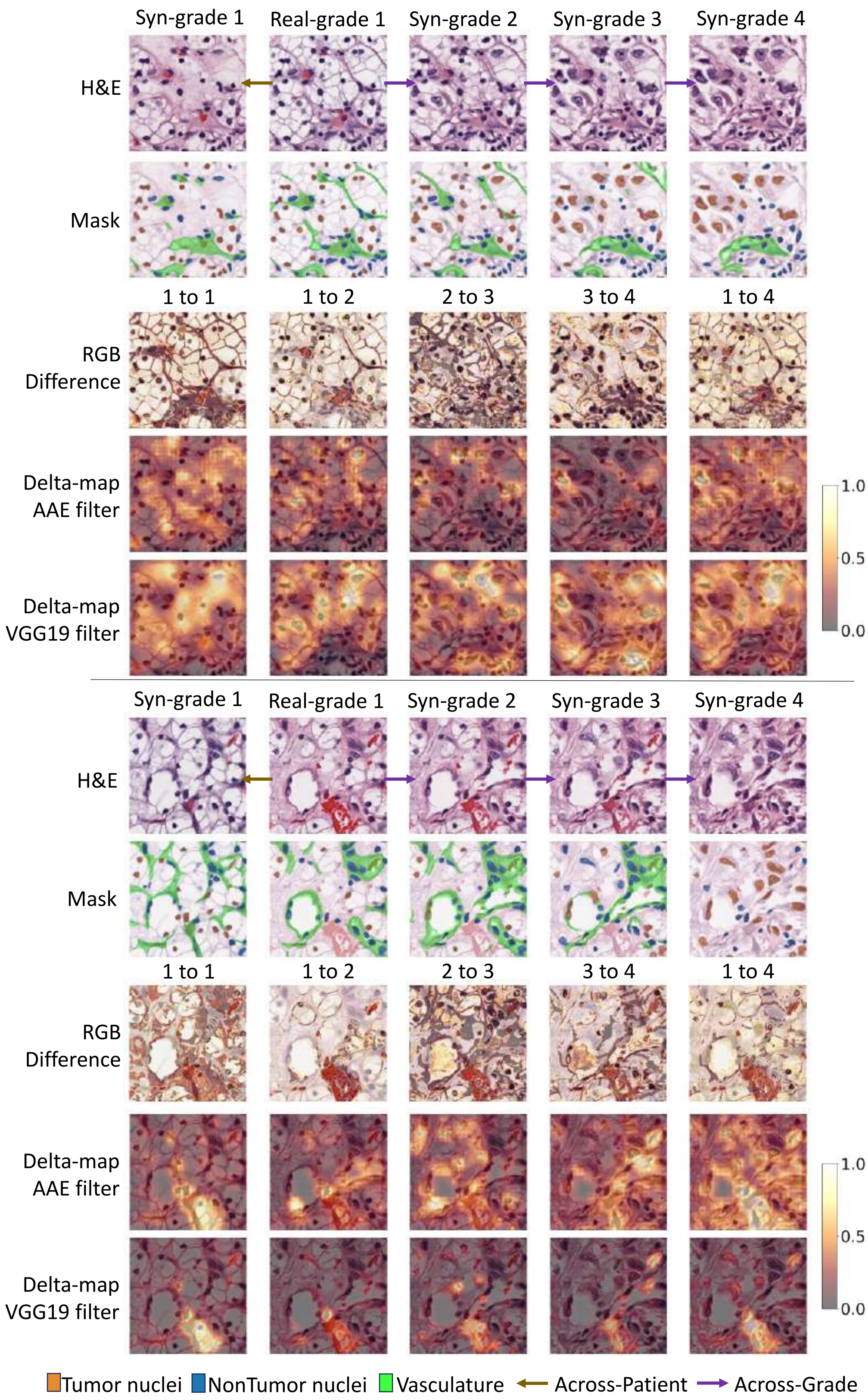
Syn-grade 1
Real-grade 1
Syn-grade 2
Syn-grade 3
Syn-grade 4
H&E
Mask
1 to 1
1 to 2
2 to 3
3 to 4
1 to 4
RGB Difference
Delta-map AAE filter
Delta-map VGG19 filter
1.0
0.5
0.0
Tumor nuclei
NonTumor nuclei
Vasculature
Across-Patient
Across-Grade

**Fig. S7. Additional examples (beyond Fig. 5B) of across-patient and across-grade transitions.** Analysis was performed as in Fig 5B, except we show the activation using a grade classification model in addition to the AAE. In the across-patient (grade 1 to grade 1) transition the resulting synthetic tissues maintain the overall spatial architecture of nuclei while exhibiting distinct tissue styles in grade 1. This underscores DIFFAE's ability to disentangle tissue-specific features and generate semantically meaningful representations. In the across-grade transition, real tissue semantic vectors are incrementally shifted toward grade 4. Intermediate points along this semantic trajectory are captured and synthesized with the original stochastic terms, generating synthetic tissues corresponding to grades 2, 3, and 4. RGB difference maps and Delta maps were each computed between the two images indicated by the column label: the across-patient pair (1 to 1), the consecutive grade steps (1 to 2, 2 to 3, 3 to 4), and the direct grade 1 to grade 4 jump (1 to 4). RGB difference is the mean absolute difference of raw RGB intensities; Delta maps were computed using CNN filters from the first 9 layers of the pretrained AAE and the first 13 layers of the VGG19 classifier.

**Fig. S8.**

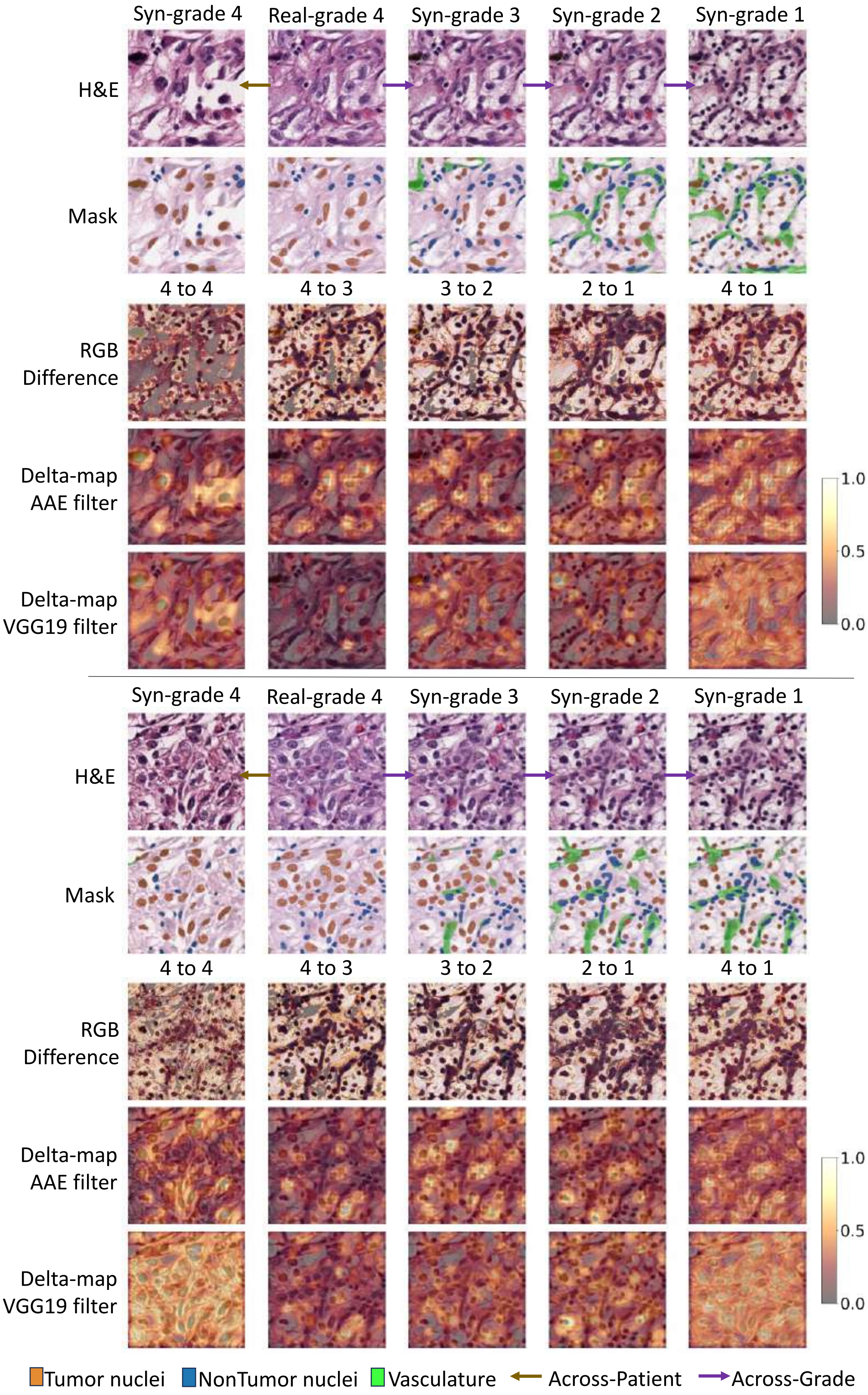

Syn-grade 4
Real-grade 4
Syn-grade 3
Syn-grade 2
Syn-grade 1
H&E
Mask
4 to 4
4 to 3
3 to 2
2 to 1
4 to 1
RGB Difference
Delta-map AAE filter
Delta-map VGG19 filter
1.0
0.5
0.0
Syn-grade 4
Real-grade 4
Syn-grade 3
Syn-grade 2
Syn-grade 1
H&E
Mask
4 to 4
4 to 3
3 to 2
2 to 1
4 to 1
RGB Difference
Delta-map AAE filter
Delta-map VGG19 filter
1.0
0.5
0.0
Tumor nuclei
NonTumor nuclei
Vasculature
Across-Patient
Across-Grade

**Fig. S8. Examples of across-patient and across-grade transitions starting from grade 4 with corresponding segmentation masks and Delta-Marches from CNN layers of the AAE model and VGG19 classifier.** This figure shows two latent transitions starting from grade 4 with similar analysis performed in Fig. S7 (and Fig. 5B). Real grade 4 tissue images from different patients were processed by DIFFAE, extracting their semantic vectors and stochastic terms. For the across-grade transition, grade 4 semantic vectors were gradually shifted toward grade 1, generating synthetic tissues for grades 3, 2, and 1. RGB difference maps and Delta maps were each computed between the two images indicated by the column label: the across-patient pair (4 to 4), the consecutive grade steps (4 to 3, 3 to 2, 2 to 1), and the direct grade 4 to grade 1 jump (4 to 1). Delta maps use CNN filters from the first 9 layers of the AAE and the first 13 layers of the VGG19 classifier. Delta-Marches heatmaps capture significant morphological changes in nuclei across grades. Additionally, differences in nuclei texture between real and synthetic tissues within grade 4 are evident.

**Fig. S9.**

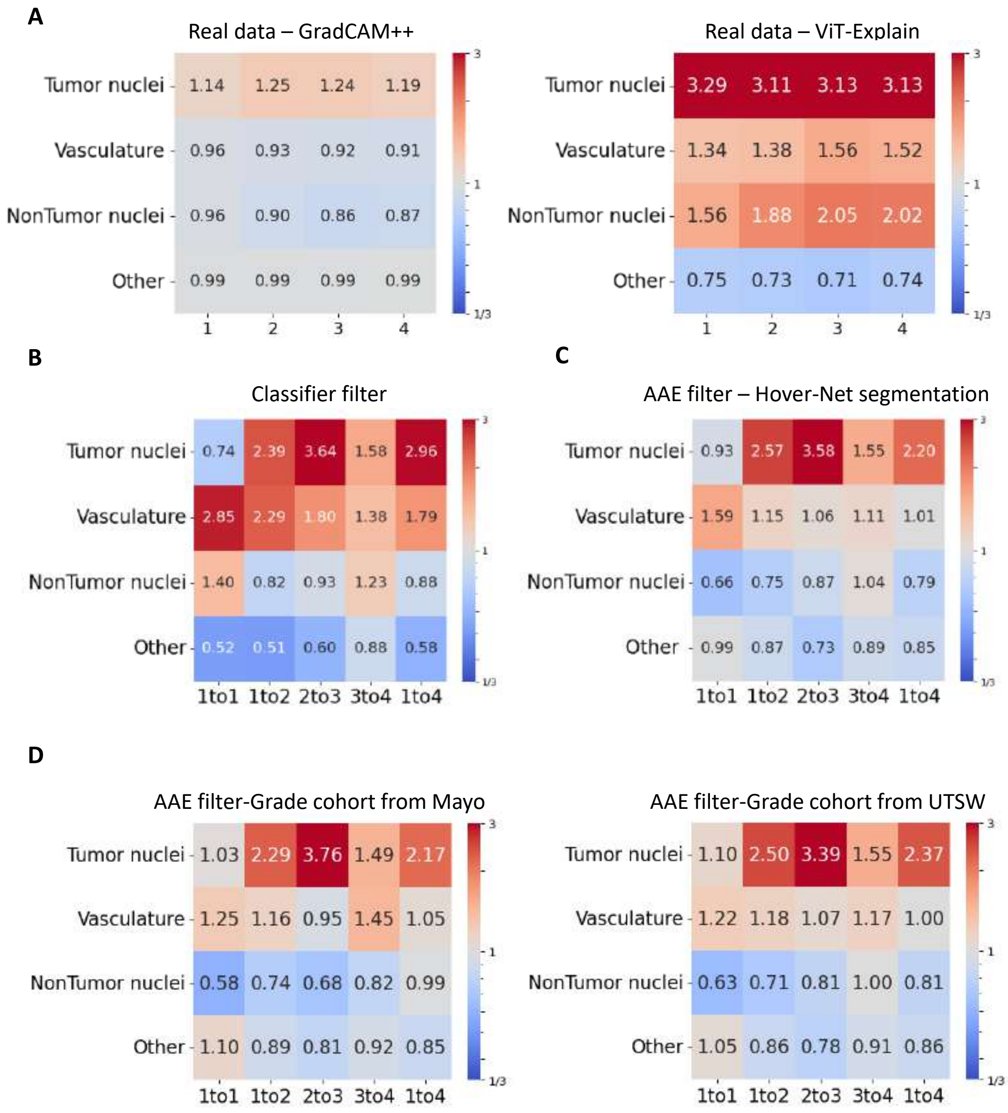


**Fig. S9. Comparison of Delta-Marches with classifier-based activation maps and evaluation of filter-model and segmentation robustness for cell-type localization.** We compared cell-type localization across different activation-generation strategies and filter models by calculating the median Jaccard similarity ratio between the binarized activation map and the corresponding cell-type mask, relative to a randomly selected mask. The activation binarization and the Jaccard ratio computation follow the same procedure as in Fig. 5C. **(A)** The median Jaccard ratio is calculated using activations obtained from Grad-CAM++ (left) and ViT-Explain (right) applied to real data, with 1,000 patches per grade. The random mask for each sample is taken from the other 999 masks having the same grade. Grad-CAM++ is applied to the VGG19 tumor grade classifier, while ViT-Explain is applied to the ViT model. Grad-CAM++ shows little enrichment in any component, whereas ViT-Explain enriches most strongly at tumor nuclei but also at non-tumor nuclei and vasculature. **(B)** Jaccard ratios for Delta-Maps generated by Delta-Marches using the first 13 layers of a VGG19-based grade classifier as the CNN filter. Synthetic grade 1-to-grade 4 transitions were analyzed (N = 977), with each sample traversing all four grades. Tumor-nuclei enrichment reproduces the pattern obtained with the AAE filter (Fig. 5C), indicating robustness to CNN filter choice. **(C)** Jaccard ratios computed using HoVer-Net-derived nuclei masks as an independent nuclei segmentation baseline. Nuclei classes are assigned by majority voting from the Rajaram lab pixel-level mask. Tumor-nuclei enrichment is preserved, showing that the result does not depend on our in-house nuclei segmentation. **(D)** Institution-stratified analysis of Delta-Marches using the AAE-based filter on the same grade 1-to-grade 4 transition cohort as in (B), showed consistent enrichment in both Mayo Clinic (N = 265) and UTSW (N = 712) samples, indicating that the result is not driven by site-specific technical variation.

**Fig. S10.**

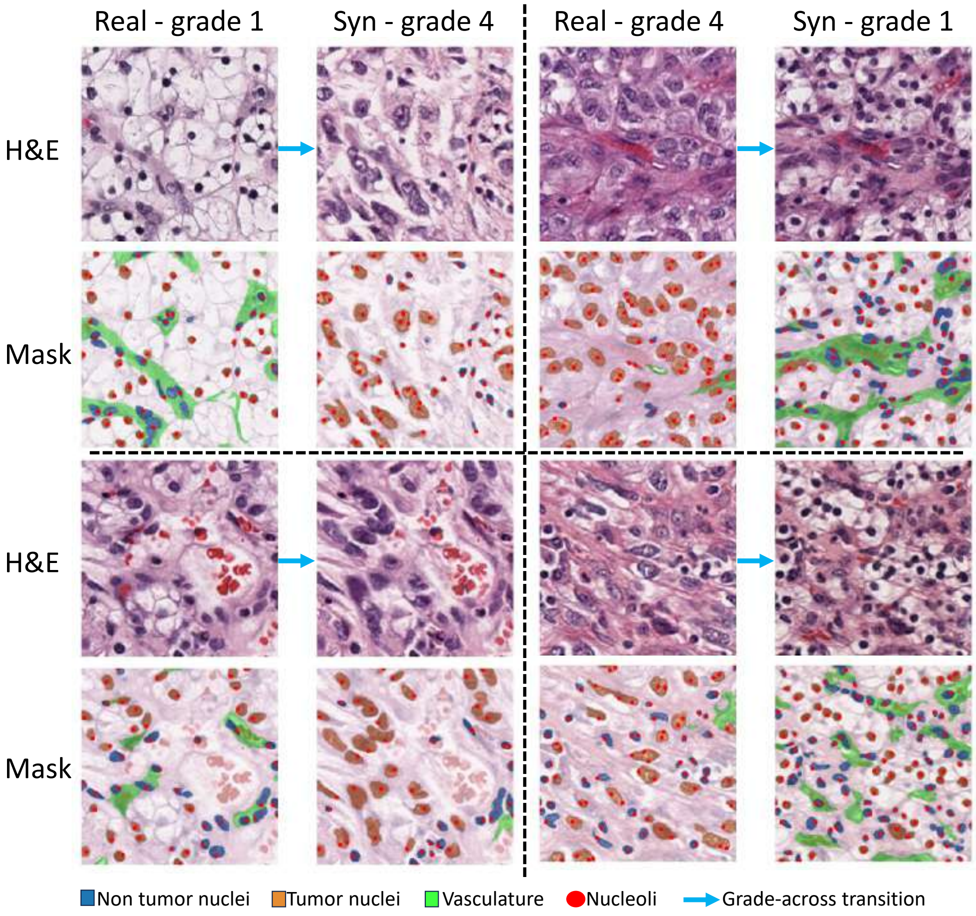


**Fig. S10. "Nucleoli" detection in Hematoxylin sections using image processing techniques.** The figure presents examples of "nucleoli" detection within nuclei from Hematoxylin sections, leveraging the HED color space and isolating the hematoxylin channel. "Nucleoli", identified as darker, separate spots, are detected as local maxima within smoothened images. Despite occasional false "nucleoli" predictions in non-tumor nuclei due to resolution limitations, the method reveals a higher average number of "nucleoli" in synthetic high-grade images compared to lower-grade images.

**Fig. S11.**

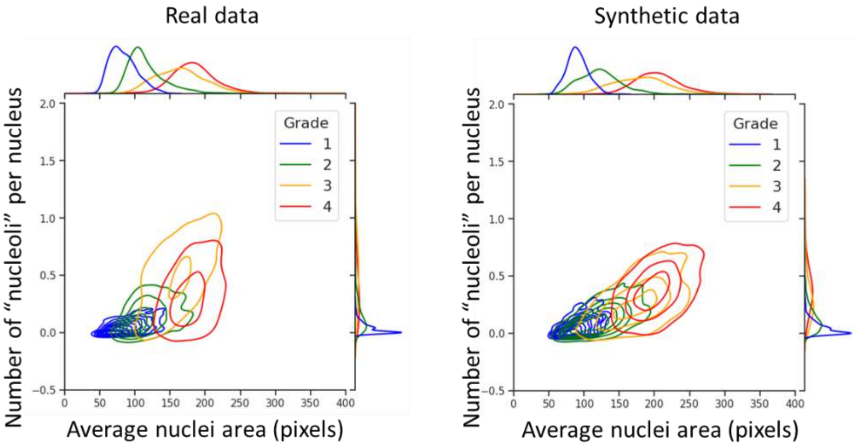


**Fig. S11. Synthetic images show greater consistency between different grade associated features than real images.** Contour plots show the correlation between tumor nuclei size and "nucleoli" number in real data and in synthetic data (N = 500 patches per grade).

**Fig. S12.**

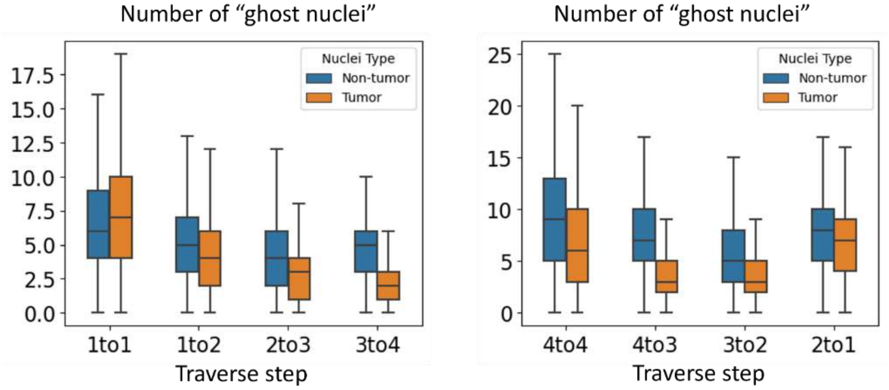


**Fig. S12: Number of "ghost nuclei" detected across grade transitions.** For each transition from grade 1 to 4 (N = 977) and from grade 4 to 1 (N = 556), we identify nuclei that appear in one segmentation mask but not the other, indicative of generative artifacts.

**Table S1.**

| Institution | Grade | Number of slides | Number of patients | Number of patches |
|---|---|---|---|---|
| Mayo | 1 | 8 | 8 | 8913 |
| | 2 | 11 | 11 | 4713 |
| | 3 | 6 | 6 | 7876 |
| | 4 | 3 | 3 | 5688 |
| UTSW | 1 | 19 | 19 | 21087 |
| | 2 | 47 | 44 | 25287 |
| | 3 | 21 | 19 | 22124 |
| | 4 | 13 | 12 | 24312 |

**Table S1. Distribution of images in the Grade cohort.** The table summarizes the number of slides, patients, and sampled patches for each tumor grade, stratified by institution. In total, 120,000 patches were used, with 30,000 patches sampled from each grade. The mean tumor grade is 2.54 for UTSW and 2.38 for Mayo.

**Table S2.**

| Grade of synthetic images | Grade of real original images | Number of images |
|---|---|---|
| 1 | 3 | 6 |
| 1 | 4 | 4 |
| 2 | 3 | 8 |
| 2 | 4 | 2 |
| 3 | 1 | 2 |
| 3 | 2 | 8 |
| 4 | 1 | 6 |
| 4 | 2 | 4 |

**Table S2. Summary of synthetic images used for pathologist evaluation.** For the assessment (N = 10 patches per grade), 20 synthetic patches representing low-grade images (grades 1 and 2) were generated by performing latent space transitions from high-grade images (grades 3 and 4) toward grade 1. Similarly, 20 synthetic high-grade patches (grades 3 and 4) were generated by transitioning from low-grade images toward grade 4.

**Table S3.**

| Class | Number of images |
|---|---|
| adenocarcinoma | 4567 |
| artifact | 4169 |
| highgrade_dysplasia | 3057 |
| inflammation | 1026 |
| lowgrade_dysplasia | 5000 |
| normal | 5000 |
| resection_edge | 541 |
| suspicious_for_invasion | 681 |
| tumor_necrosis | 624 |

**Table S3. Summary of images in Colorectal data used for training DIFFAE.**
